# GCP-VQVAE: A Geometry-Complete Language for Protein 3D Structure

**DOI:** 10.1101/2025.10.01.679833

**Authors:** Mahdi Pourmirzaei, Alex Morehead, Farzaneh Esmaili, Jarett Ren, Mohammadreza Pourmirzaei, Dong Xu

## Abstract

Converting protein tertiary structure into discrete tokens via vector-quantized variational autoencoders (VQ-VAEs) creates a language of 3D geometry and provides a natural interface between sequence and structure models. While pose invariance is commonly enforced, retaining chirality and directional cues without sacrificing reconstruction accuracy remains challenging. In this paper, we introduce GCP-VQVAE, a geometry-complete tokenizer built around a strictly SE(3)-equivariant GCPNet encoder that preserves orientation and chirality of protein backbones. We vector-quantize rotation/translation-invariant readouts that retain chirality into a 4 096-token vocabulary, and a transformer decoder maps tokens back to backbone coordinates via a 6D rotation head trained with SE(3)-invariant objectives.

Building on these properties, we train GCP-VQVAE on a corpus of 24 million monomer protein backbone structures gathered from the AlphaFold Protein Structure Database. On the CAMEO2024, CASP15, and CASP16 evaluation datasets, the model achieves backbone RMSDs of 0.4377 Å, 0.5293 Å, and 0.7567 Å, respectively, and achieves 100% codebook utilization on a held-out validation set, substantially outperforming prior VQ-VAE–based tokenizers and achieving state-of-the-art performance. Beyond these benchmarks, on a zero-shot set of 1 938 completely new experimental structures, GCP-VQVAE attains a backbone RMSD of 0.8193 Å and a TM-score of 0.9673, demonstrating robust generalization to unseen proteins. Lastly, we show that the Large and Lite variants of GCP-VQVAE are substantially faster than the previous SOTA (AIDO), reaching up to ∼ 408 × and ∼ 530 × lower end-to-end latency, while remaining robust to structural noise. We make the GCP-VQVAE source code, zero-shot dataset, and its pretrained weights fully open for the research community: https://github.com/mahdip72/vq_encoder_decoder

## 1 Introduction

Proteins are the molecular machines of life, and their function is intricately tied to their three-dimensional structures (Bertoline et al., 2023; Tripathi et al., 2025). Understanding and predicting these structures remains one of the central challenges in computational biology (Jänes & Beltrao, 2024). Just as natural language is governed by grammatical and contextual rules, protein 3D structures exhibit spatial patterns and constraints that suggest an underlying “grammar” of folds and interactions (Ruff et al., 2022; Kilgore et al., 2025; Weissenow & Rost, 2025).

Despite advances in understanding and prediction of protein structures, effective representation of the 3D geometry of proteins in a form easily suitable for generative modeling remains an open problem (Draizen et al., 2024; Lu et al., 2025). While recent methods have begun to leverage generative AI, such as diffusion models and autoregressive frameworks, to produce full-atom structures or backbone coordinates, they still lag behind other domains such as language or vision in terms of reconstruction precision, scalability to large and diverse datasets, and openness for the broader research community, like natural language or image and video generation (Yuan et al., 2025b). These gaps continue to constrain our ability to build powerful, general-purpose protein models using modern AI techniques.

Beyond generative modeling, learning a discrete language for protein 3D structure opens up a wide range of downstream applications. First, compressing 3D coordinates into compact sequences of integer codes—while preserving accurate reconstruction—can substantially reduce storage and transmission costs for structural data (Kim et al., 2023). Second, discrete structural representations enable fast, alignment-style comparison of protein shapes, analogous to multiple sequence alignment in sequence space (Van Kempen et al., 2024). Third, in learnable quantization frameworks such as vector-quantized autoencoders, these codes can be decoded into semantically rich continuous embeddings (Yuan et al., 2025b), facilitating structure-aware feature extraction for classification, clustering, structure-based comparison/search, and other predictive tasks. Finally, unifying protein sequence and structure through a shared discrete representation may pave the way for multimodal generative models that bridge amino acid sequences and 3D folds within a common language modeling framework (Hsieh et al., 2025).

However, most existing protein-structure VQVAEs are either closed-source or only partially released (e.g., code without full evaluation scripts or strongest checkpoints), which impedes reproducible comparison (Gao et al., 2024c; Hayes et al., 2025). Furthermore, the publicly available baselines often generalize weakly to *unseen* proteins. Consequently, the field lacks a fully opensource, high-accuracy tokenizer with transparent training and evaluation that demonstrably transfers to new proteins.

### Contributions

(1) We introduce GCP-VQVAE, a geometry-complete tokenizer that preserves orientation and chirality while producing pose-invariant codes that support missing coordinates. (2) We scale training to 24M monomers and report exhaustive evaluations (CAMEO2024, CASP14/15/16) and a zero-shot suite of newly deposited experimental structures. (3) Our large model achieves state-of-the-art reconstruction on a diverse range of unseen 3D structures, and stays ahead of other open-source methods. On the zero-shot set, GCP-VQVAE attains a backbone RMSD of 0.8193 Å and a TM-score of 0.9673. (4) Building on Yuan et al. (2025b), we evaluate our continuous protein-structure representations using probes of varying complexity, quantifying how readily task-relevant information can be extracted for supervised downstream tasks. (5) We evaluate practical deployment properties of GCP-VQVAE by benchmarking end-to-end inference latency across sequence lengths and quantifying reconstruction robustness under structural noise and partial-structure inputs. (6) We open-source our implementation, trained checkpoints, and datasets to support reproducible bench-marking by the community.

## 2 Related Work

An early and influential approach to casting protein 3D structure as a discrete language is FoldSeek van Kempen et al. (2022), which learns a 20-state 3Di alphabet with a VQ-VAE trained for evolutionary conservation and encodes structures as token sequences for ultra-fast k-mer–based local/-global alignment. Building on the notion of a discrete structural language—but targeting generative reconstruction rather than search—the FoldToken series (Gao et al., 2025; 2024a;b;c) develops a VQ-VAE–style tokenizer and decoder: FoldToken introduces a SoftCVQ fold language with joint sequence–structure generation; FoldToken2 stabilizes quantization and extends to multi-chain settings; FoldToken3 mitigates gradient/class-space issues to reach 256-token compression with minimal loss; and FoldToken4 unifies cross-scale consistency and hierarchies in a single model, reducing redundant multi-scale training and code storage.

Yuan et al. (2025b) introduces AminoAseed codebook reparameterization plus Pareto-optimal K ×D sizing, proposes structtokenbench—a fine-grained evaluation suite—and diagnoses codebook underutilization in VQVAE PSTs. Concurrently, Gaujac et al. (2024) tokenizes protein backbones with a VQ autoencoder (codebooks 4k–64k); its main open-source limitation is that it does not support sequences shorter than 50 or longer than 512 amino acids.

ESM-3 (Hayes et al., 2025) couples its multimodal transformer with a VQVAE structure tokenizer that discretizes local 3D geometry into structure tokens; structure, sequence, and function are jointly trained under a masked-token objective. The tokenizer uses an SE(3)-aware module within the encoder, and ESM-3 employs a 4 096-code structure codebook (plus special tokens) for downstream generation and masked reconstruction.

Across prior structure tokenizers, neither openness nor accuracy are yet satisfactory. The FoldToken line spans four variants, but to our knowledge, only FoldToken-4 offers a partial open release, without the strongest checkpoints, limiting reproducibility (Gao et al., 2024c). ESM-3 exposes an internal VQ-VAE tokenizer, yet public artifacts lack fully documented training/evaluation procedures and best weights, making directly comparable reconstruction benchmarking difficult (Hayes et al., 2025). The open VQ autoencoder of Gaujac et al. (2024) supports a restricted length window (e.g., ∼ 50–512 residues), precluding fair assessment on long chains where reconstruction accuracy degradation is most evident. Finally, FoldSeek’s learned 3Di alphabet targets ultra-fast structure search and does not provide a generative decoder from discrete codes back to coordinates, so reconstruction fidelity cannot be evaluated (van Kempen et al., 2022). Consequently, the community still lacks a fully open, end-to-end tokenizer with released best weights and source codes that attains high reconstruction accuracy on *unseen* proteins.

## 3 Dataset

We began with the latest release of UniRef50 (Consortium, 2019) ^1^, which clusters protein sequences at 50% identity and thus offers a natural, non-redundant scaffold for large-scale structure modeling. For every UniRef50 entry with a corresponding model in the AlphaFold Database (AFDB; Varadi et al. (2022)), we downloaded the per-protein structure (PDB format at collection time). Limiting to UniRef50 reduces near-duplicate leakage by ensuring that homologs within clusters do not exceed 50% identity. After parsing and splitting multi-chain records into individual chains, this procedure yielded approximately 42M single-chain PDB samples.

From this AFDB ∩ UniRef50 pool, we drew a uniform random sample of 24M single-chain structures to form the training set. This down-sampling keeps training throughput tractable while preserving the global distribution of lengths, folds, and taxa present in the full pool. To support our lightweight model variant, we also employed the `afdb_rep_v4` dataset (Jamasb et al., 2024), which contains approximately 2.2 million cluster-representative structures, serving as a data-efficient alternative to the full training corpus.

We evaluate on three held-out test suites capturing complementary shifts (see Table 1): *A* low-confidence AF2 SwissProt models, *B* species-diverse (97-species) taxonomic shift, and *C* high-confidence structures from under-represented taxa. All suites are per-chain, length-filtered, deduplicated, and NaN-aware; a balanced validation set of 6 000 chains (2k per suite) is sampled disjointly. Full curation criteria, thresholds, and counts are provided in Appendix B.

**Table 1:** Dataset and benchmark statistics. Training data is deduplicated at 100% sequence identity against all validation/test splits and external benchmarks (CAMEO2024, CASP14–16). We also include a zero-shot set of 1 938 newly deposited experimental monomer chains (PDB, 2024–2025).

| Split | Source | Selection criterion | # Samples |
| --- | --- | --- | --- |
| Training (Large) | AFDB ∩ UniRef50 | Uniform random sample from ∼42M pool | 24 000 000 |
| Training (Lite) | AFDB | Cluster representatives (afdb_rep_v4) | 2 200 000 |
| Validation | From Tests A/B/C | 2k random per test (A,B,C) merged | 6 000 |
| Test set A | PDB DB | cross-referenced AF pLDDT $\leq 70$; len $\geq 25$; $< 25$ consecutive missing residues; dedup | 31 112 |
| Test set B | AFDB | 97 species; ∼2.5k/species; len $\geq 25$; dedup | 17 353 |
| Test set C | PDB DB | under-represented taxonomies in SwissProt; pLDDT $\geq 95$; $< 25$ consecutive missing residues; dedup | 31 867 |
| Zero-shot (experimental) | PDB DB | recent experimental monomers (2024–2025); len 25–2048; dedup; $< 50\%$ seq id vs train | 1 938 |
| CAMEO2024 | CAMEO | - | 574 |
| CASP14 | CASP | - | 33 |
| CASP15 | CASP | - | 45 |
| CASP16 | AF3 predictions | - | 104 |

To probe generalization to genuinely new structures, we assembled a zero-shot suite from *recently deposited experimental* monomeric chains in the PDB (calendar years 2024–2025). Multi-chain entries were split into per-chain backbones, chains shorter than 25 residues or longer than 2 048 residues were discarded, and we retained entries even when they contained missing coordinates (NaN) to reflect typical experimental gaps, yielding 76 503 pdbs in total. We deduplicated using structural clustering with FoldSeek easy-cluster (alignment type 2, tmscore threshold 0.80, cov-mode 0, lddt-threshold 0.65), retaining one representative per cluster. This resulted in 10 689 clusters (van Kempen et al., 2022). For leakage control, each representative was compared with foldseek `easy-search` against (i) the AFDB ∩ UniRef50 set used for our GCP-VQVAE Large model training and (ii) the `afdb_rep_v4` (Barrio-Hernandez et al., 2023) representative set used to pre-train the GCPNet encoder in ProteinWorkshop (Jamasb et al., 2024) and training the GCP-VQVAE Lite model. We *excluded* any candidate with ≥ 50% structure similarity to any chain in those sources; the resulting zero-shot set comprises 1 938 monomer chains.

### Independent benchmarks

In addition to the internal splits, we evaluate on community benchmarks to facilitate comparison with prior work: CAMEO2024 (Robin et al., 2021; Leemann et al., 2023), CASP14 (Kryshtafovych et al., 2021), CASP15 (Kryshtafovych et al., 2023), and AlphaFold3 (Abramson et al., 2024) predictions for CASP16 (Yuan et al., 2025a) targets. Each benchmark is processed with the same per-chain extraction and deduplication pipeline and is used strictly for out-of-distribution evaluation.

Table 1 summarizes the composition and selection criteria of all splits. Counts after converting multi-chain inputs into per-chain samples. Also, all samples are truncated to a maximum of 2 048 amino acids in length.

## 4 GCP-VQVAE Architecture

The proposed model couples (i) a geometry-complete SE(3)-equivariant GCPNet encoder that maps protein backbone coordinates (N–C_*α*_–C) to per-residue embeddings, with (ii) a transformer-based VQVAE that discretizes these embeddings into pose-invariant code indices and reconstructs the backbone in 3D. An overview is shown in Figure 1B; full architectural details are provided in Appendix A.

**Figure 1.**
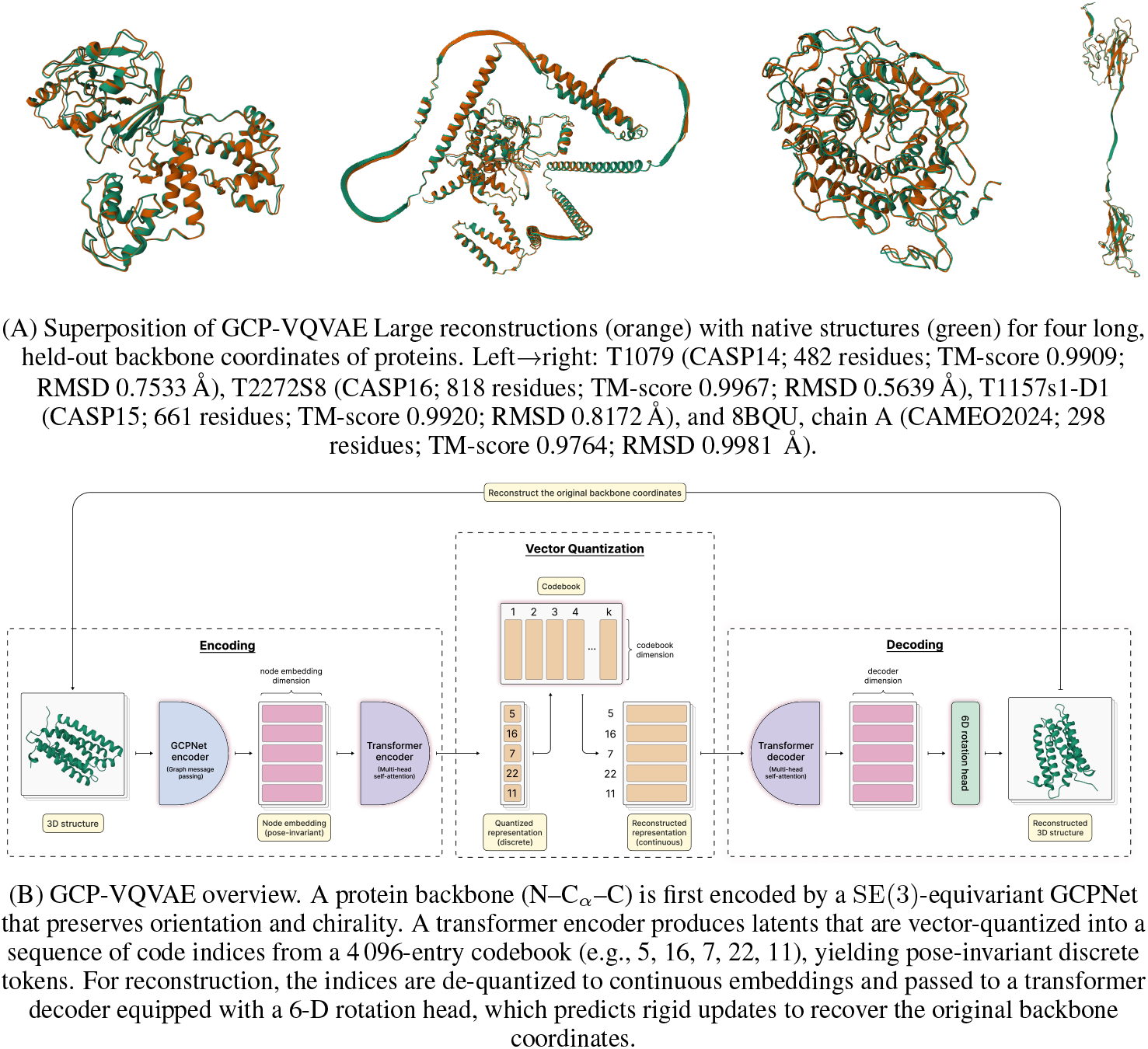
Qualitative reconstructions (A) and an overview of the GCP-VQVAE architecture (B).

### 4.1 GCPNet Encoder

GCPNet (Morehead & Cheng, 2024b) performs scalar–vector message passing while explicitly preserving orientation and chirality via local edge frames, yielding SE(3)-equivariant backbone representations. We use its final node embeddings as the continuous input sequence to the VQVAE (Appendix A.1).

### 4.2 VQVAE

The VQVAE follows the standard *encoder* → *vector* → *quantization decoder* pipeline: a transformer encoder produces latent vectors **z**, which are mapped to the nearest codebook entry **e**_*k*_ (discrete index *k*), and a transformer decoder maps the de-quantized embeddings back to structure. To generate coordinates, the decoder is equipped with a 6D rotation head (Zhou et al., 2019) that predicts rigid updates and applies them to a fixed local backbone template (Appendix A.2).

### 4.3 Training objective

We supervise reconstruction with a weighted sum of pose-invariant geometric losses. We first align the ground-truth backbone to the prediction using Kabsch alignment (proper rotation + translation; no reflections), and then compute an aligned MSE term, a backbone distance-matrix loss, and a pose-invariant *relative* direction/orientation loss. The latter matches internal backbone directions and plane normals by comparing their pairwise dot products, and is therefore invariant to global rigid transformations:

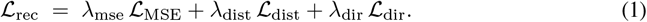

The full objective adds standard VQ codebook and commitment losses (Van Den Oord et al., 2017), regularized to encourage diverse code usage and orthogonality:

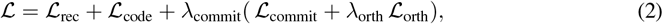

with implementation details deferred to Appendix A.2.

## 5 Experiments

In this section, optimization is used with AdamW (Loshchilov, 2017) and a cosine-annealed learning-rate schedule with warm-up steps (Loshchilov & Hutter, 2016). Experiments were implemented in PyTorch 2.7 (Ansel et al., 2024) ^2^ with mixed-precision (BF16) training (Kalamkar et al., 2019; Micikevicius et al., 2017) on four nodes of NVIDIA 8 × A100 GPUs. For the GCPNet encoder, we initialized from the ProteinWorkshop checkpoint (Jamasb et al., 2024). Both light and large GCP-VQVAE configurations and exact hyperparameter values are listed in Appendix C; Table 9 and 10.

For the GCP-VQVAE Lite variant, we employed the afdb_rep_v4 dataset from the ProteinWork-shop benchmark (Jamasb et al., 2024), which consists of approximately 2.27 million representative AlphaFold structures. This roughly 10-fold reduction in training data relative to our primary model allows us to demonstrate the architectural efficiency and robust performance of GCP-VQVAE even under significantly reduced computational budgets.

Training of the large model proceeded in two stages on the same training split. In Stage 1, sequences are truncated to 512 residues. In Stage 2, the maximum length is increased to 1 280 residues. In Stage 2, we also introduce a simple NaN-masking augmentation to handle missing coordinates; implementation details are provided in Appendix C (Figure 6). At the end of Stage 2 training, on the validation set, the model attains MAE 0.2239, RMSD 0.4281, GDT-TS 0.9856, and TM-score 0.9889 with 100% codebook utilization. Similarly, the training of the lite model proceeded by one stage only with 1 280 residues. On the validation set, the model attains MAE 0.3961, RMSD 0.5707, GDT-TS 0.9327, and TM-score 0.9741 with 100% codebook utilization. Per-test-set result of the large model appears in Table 3.

Figure 2 compares end-to-end inference latency across sequence lengths measured on a Nvidia A6000 Ada GPU, showing that GCP-VQVAE scales efficiently to long proteins relative to open baselines; full benchmarking details are provided in Appendix C.2.

**Figure 2.**
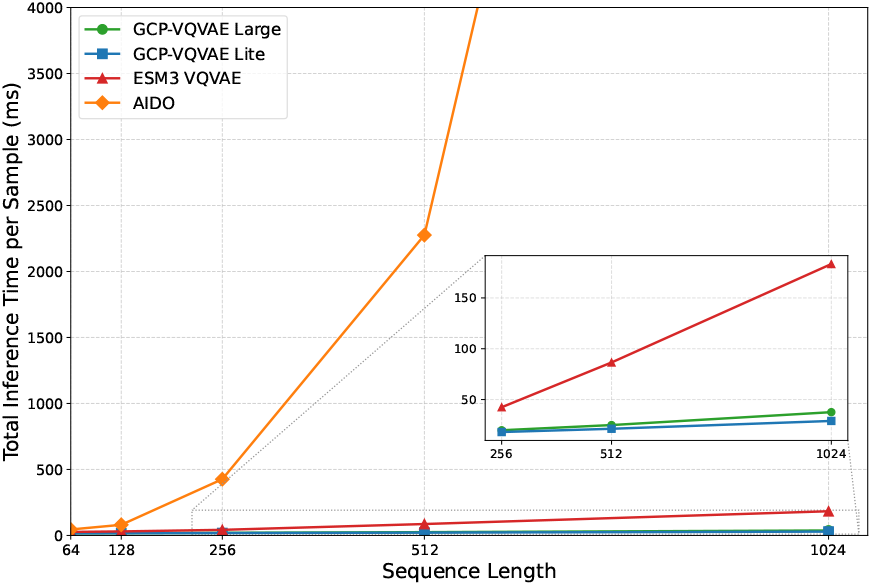
End-to-end inference time per protein (ms) as a function of sequence length for GCP-VQVAE (Large/Lite) and top-performing open baselines (ESM3 VQVAE, AIDO). The inset highlights the long-sequence regime (256–1024 residues).

To quantify the contribution of each decoder reconstruction term, we performed a small ablation study over the aligned MSE (ℒ _MSE_), distance-matrix (ℒ _dist_), and direction/orientation (ℒ _dir_) losses. All ablations were trained with the GCP-VQVAE Lite architecture on 100k randomly sampled training structures for 32 epochs, using a maximum sequence length of 512 residues. Table 4 reports reconstruction quality under different loss combinations.

We compared against open-source structure tokenizers. Among them, in practice, ESM-3 does not provide documented usage or official evaluation scripts for its VQ-VAE; we therefore reproduced its evaluation using the publicly released weights and their GitHub codebase. The original FoldToken-4 evaluation repository was prohibitively slow, so we re-implemented it, yielding a ∼ 20 × speed-up with negligible loss in reconstruction rate. For all other baselines, we relied on the officially released implementations and corresponding checkpoints. See Table 2 for aggregate metrics; Figure 5 visualizes the full RMSD error across test sets.

**Table 2:** Reconstruction comparison with the available open-source structure tokenizer methods. FoldToken-4, AminoAseed, and DPLM-2 use 256, 512, and 8 192 vocab sizes, respectively. Other methods use 4 096 vocab size. The Structure Tokenizer of Gaujac et al. (2024) only supports sequences of length 50–512. For AIDO, sequences longer than 1 024 are omitted due to OOM.

| Dataset | Metric | GCP-VQVAE<br>Large (Ours) | GCP-VQVAE<br>Lite (Ours) | AIDO<br>(Zhang et al., 2024) | ESM-3 VQVAE<br>(Hayes et al., 2025) | DPLM-2 VQVAE<br>(Wang et al., 2024b) | FoldToken 4<br>(Gao et al., 2024c) | (Gaujac et al., 2024) | AminoAseed<br>(Yuan et al., 2025b) |
| --- | --- | --- | --- | --- | --- | --- | --- | --- | --- |
| CASP14 | TM-score ↑ | **0.9890** | 0.9751 | <u>0.9831</u> | 0.9734 | 0.9056 | 0.8989 | 0.8940 | 0.3951 |
| | RMSD ↓ | **0.5431** | 0.8435 | <u>0.6587</u> | 1.0968 | 2.0896 | 2.0027 | 1.8913 | 14.6739 |
| CASP15 | TM-score ↑ | **0.9884** | 0.9665 | <u>0.9660</u> | 0.9418 | 0.8268 | 0.9136 | 0.7911 | 0.3831 |
| | RMSD ↓ | **0.5293** | 0.9219 | <u>0.8611</u> | 1.3407 | 6.3826 | 1.6856 | 8.5592 | 18.5827 |
| CASP16 | TM-score ↑ | **0.9857** | 0.9757 | 0.8805 | 0.9103 | 0.7852 | 0.8211 | 0.8345 | 0.3433 |
| | RMSD ↓ | **0.7567** | <u>1.0614</u> | 4.9773 | 4.4918 | 14.1934 | 5.4115 | 6.9375 | 25.9508 |
| CAMEO2024 | TM-score ↑ | **0.9918** | 0.9794 | <u>0.9851</u> | 0.9736 | 0.9243 | 0.9346 | 0.9114 | 0.4632 |
| | RMSD ↓ | **0.4377** | 0.7401 | <u>0.5867</u> | 1.0156 | 1.941 | 1.4069 | 2.1726 | 12.6203 |
| Zero-Shot | TM-score ↑ | **0.9673** | 0.9466 | 0.9202 | 0.8466 | 0.7137 | 0.8256 | 0.7662 | 0.4155 |
| | RMSD ↓ | **0.8193** | <u>1.1152</u> | 2.0105 | 6.5075 | 12.2182 | 4.5465 | 10.4405 | 19.2612 |
| Average | TM-score ↑ | **0.9844** | 0.9687 | 0.9470 | 0.9291 | 0.8311 | 0.8788 | 0.8394 | 0.4000 |
| | RMSD ↓ | **0.6172** | <u>0.9364</u> | 1.8189 | 2.8905 | 7.3650 | 3.0106 | 6.0002 | 18.2178 |

**Table 3:** Evaluate GCP-VQVAE large on the predefined test sets.

| Method | Test set A | Test set B | Test set C |
| --- | --- | --- | --- |
| RMSD ↓ | 0.5124 | 0.4027 | 0.4787 |
| TM-score ↑ | 0.9823 | 0.9951 | 0.9885 |

**Table 4:**
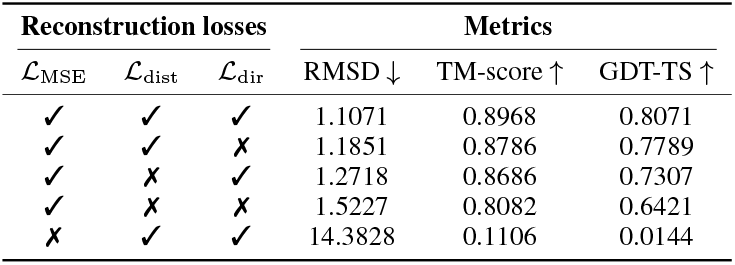
Ablation of reconstruction losses and their impact on backbone reconstruction quality.

We examined how reconstruction error varies with sequence length on the validation set (Appendix C; Figure 7). Errors increase only mildly with length, indicating stable accuracy up to 1 280 residues with only a slight variance rise for very long chains. On the zero-shot set we obtain mean RMSD 0.8193 Å and TM-score 0.9673 (Table 2), with group-wise trends summarized in Table 5; extended zero-shot analyses (Figure 9 to 15) appear in Appendix C.1.

**Table 5:** Metrics by structure group on the zero-shot dataset with respect to the GCP-VQVAE Large. Groups are defined by secondary-structure composition thresholds and are not mutually exclusive. Thresholds of each group are described in Appendix B

| Group | Count | RMSD ↓ | TM-score ↑ |
| --- | --- | --- | --- |
| All rows | 1938 | 0.8193 | 0.9673 |
| A (mainly $\alpha$) | 776 | 0.7537 | 0.9707 |
| B (mainly $\beta$) | 309 | 0.7635 | 0.9762 |
| AB (mixed) | 71 | 0.6166 | 0.9878 |
| D (coil-rich/disordered) | 552 | 0.9476 | 0.9498 |
| No-NaN residues | 1231 | 0.6179 | 0.9755 |
| Has-NaN residues ($\geq 1$) | 707 | 1.1699 | 0.9531 |

We additionally evaluate reconstruction robustness under coordinate perturbations and partial-structure inputs, reporting ΔRMSD degradation relative to clean reconstructions in Appendix C.3.

### 5.1 Codebook Evaluation

We analyzed the distribution and informational properties of the learned discrete codes on the zero-shot test set (Table 6). GCP-VQVAE demonstrates superior codebook efficiency, achieving 100% utilization of its vocabulary. In terms of structural grammar, the Large model exhibits a transition from high unigram entropy (*H*_1_ = 11.3 bits) to extremely low conditional trigram entropy (≈ 0.23 bits). This sharp drop implies that the model has learned a highly structured *syntax* for protein co-ordinates, where local dependencies (likely corresponding to secondary structure constraints) make token sequences highly predictable locally, similar to DPLM-2 and ESM3 models.

**Table 6:**
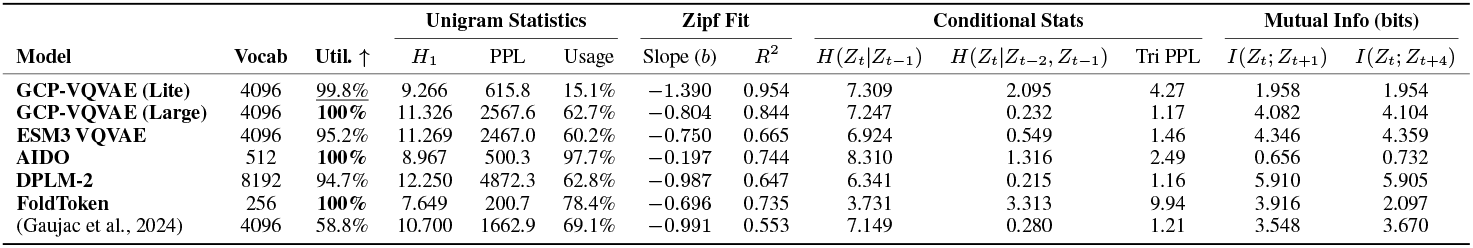
Comprehensive codebook analysis comparing structure tokenization methods on the zero-shot test set.

Geometrically, the reconstructed backbone dihedrals closely imitate the original distribution, validating the decoder’s precision. However, the observation (Figure 3) that individual top codes often span multiple Ramachandran regions suggests that tokens are not uniquely tied to narrow geometries but cover broader conformational modes, likely relying on sequence context for disambiguation (Appendix C.5).

**Figure 3.**
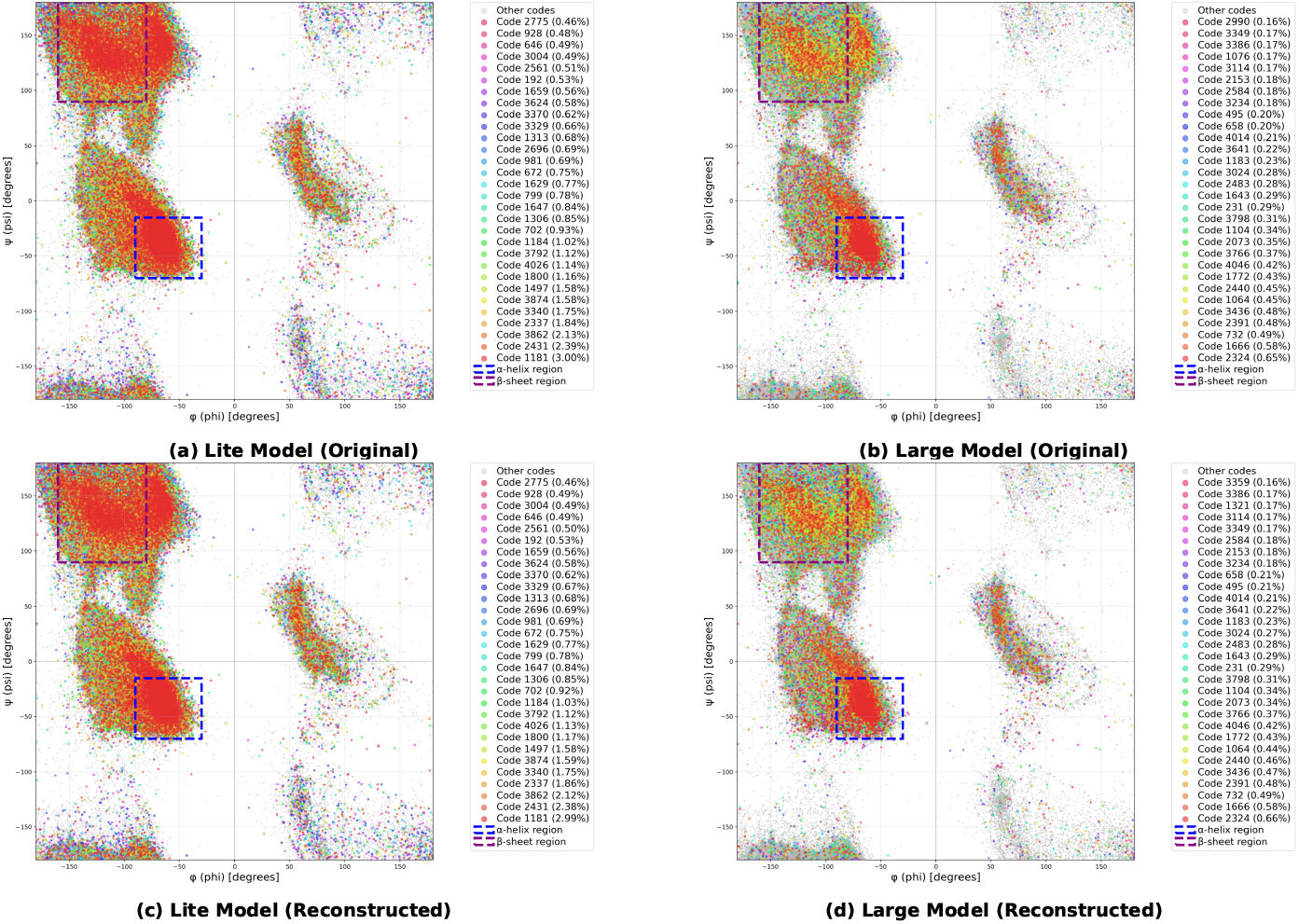
Ramachandran plot analysis of VQ codes on a zero-shot test set, colored by the top-30 most frequent codes with respect to the reconstructed and original coordinates. Dashed boxes indicate *α*-helix (blue) and *β*-sheet (purple) regions. **(a-b)** Lite model. **(c-d)** Large model. The top row shows original coordinates; the bottom row shows reconstructed coordinates.

### 5.2 Representation Evaluation

Beyond coordinate reconstruction, we evaluate the semantic quality of the learned latent space using PST benchmark (Yuan et al., 2025b). Following PST, we extract continuous (de-quantized) embeddings from our codebook and assess how well they support downstream prediction under frozen encoder settings. Since the measured utility of a representation can depend on the capacity of the evaluation head, we report results in Table 7 using three probe families: the PST-standard two-layer MLP, a stronger four-layer MLP probe, and a lightweight context-aware probe consisting of a single transformer block with the same MLP dimension.

**Table 7:** Protein Structure Tokenization benchmark across supervised Functional Site Prediction tasks. We report the two additional probes in parentheses (four-layer MLP, single-block transformer).

| VQ-VAE-based PST |  |  |  |  |  |  |  | Ours |  |
| --- | --- | --- | --- | --- | --- | --- | --- | --- | --- |
| Task | Split | Fold<br>Seek | Pro<br>Tokens | VanillaVQ | AIDO | ESM3 | AminoAseed | Our Model<br>(Large) | Our Model<br>(Lite) |
| Functional Site Prediction (AUROC% ↑) | | | | | | | | | |
| BindInt | Fold | <u>53.18</u> | 44.66 | 47.25 | 44.66 | 44.30 | 47.11 | 52.61 (50.26, 52.78) | 49.43 (49.0, **55.03**) |
|  | SupFam | 46.20 | 86.05 | 86.71 | 84.21 | **90.77** | <u>90.53</u> | 58.04 (70.61, 80.67) | 62.27 (70.27, 80.77) |
| BindBio | Fold | 52.37 | 58.47 | 62.02 | 65.50 | 62.84 | 65.73 | 75.05 (92.74, 84.01) | 80.93 (**94.61**, 91.74) |
|  | SupFam | 52.41 | 60.47 | 62.92 | 66.70 | 65.22 | 68.30 | 75.64 (93.4, 86.62) | 81.72 (**94.08**, 90.08) |
| BindShake | Org | 53.40 | 59.82 | 67.04 | 69.28 | 66.10 | <u>69.61</u> | 66.97 (67.96, 72.71) | 66.34 (67.11, 68.08) |
| CatInt | Fold | 53.43 | 58.16 | 58.89 | 57.30 | <u>61.09</u> | **62.19** | 55.92 (53.15, 60.67) | 54.36 (55.49, 58.09) |
|  | SupFam | 51.41 | 83.85 | 85.00 | 81.94 | <u>89.82</u> | **91.91** | 63.22 (73.43, 84) | 62.24 (68.88, 85.18) |
| CatBio | Fold | 56.37 | 56.14 | 67.58 | 73.72 | 65.33 | 65.95 | 67.12 (79.66, 71.64) | 67.25 (**92.28**, <u>86.11</u>) |
|  | SupFam | 53.78 | 64.05 | 70.92 | 78.66 | 74.65 | 87.59 | 73.77 (<u>89.64</u>, 86.95) | 75.57 (**95.72**, 93.38) |
| Con | Fold | 49.26 | 56.23 | 56.98 | 56.64 | 55.22 | 57.23 | 52.64 (53.29, **62.97**) | 51.15 (51.56, 61.52) |
|  | SupFam | 51.39 | 74.33 | 74.60 | 73.79 | 80.53 | **86.60** | 61.31 (68.84, 84.05) | 58.66 (65.23, <u>85.79</u>) |
| Rep | Fold | 47.70 | **77.25** | <u>75.99</u> | 77.69 | 74.70 | 74.97 | 51.19 (50.95, 66.19) | 52.04 (52.79, 61.22) |
|  | SupFam | 52.53 | 78.90 | 82.09 | 78.08 | 82.36 | 84.57 | 54.17 (65.26, <u>85.77</u>) | 56.5 (54.65, **85.94**) |
| Ept | Fold | 54.52 | 54.69 | 59.28 | 60.26 | **63.69** | <u>62.16</u> | 54.24 (54.21, 58.19) | 53.05 (54.63, 56.89) |
|  | SupFam | 50.56 | <u>67.52</u> | 67.24 | 72.30 | 61.97 | **72.02** | 55.91 (58.03, 66.31) | 56.13 (58.25, 64.09) |
| Average AUROC% |  | 51.90 | 65.37 | 68.30 | 69.38 | 69.24 | 72.43 | 61.19 (68.09, <u>73.57</u>) | 61.84 (68.3, **74.92**) |
| Physicochemical Property Prediction (Spearman's $\rho$% ↑) | | | | | | | | | |
| FlexRMSF | Fold | 15.35 | 13.81 | 44.22 | 33.15 | <u>44.53</u> | **44.63** | 18.05 (20.04, 24.37) | 24.71 (29.08, 23.79) |
|  | SupFam | 11.99 | 7.62 | 39.08 | 26.93 | <u>39.68</u> | **40.99** | 20.37 (21.21, 38.89) | 25.67 (28.03, 30.52) |
| FlexBFactor | Fold | 4.17 | 6.67 | 22.32 | 18.88 | <u>23.60</u> | 21.30 | 13.66 (14.88, 18.87) | 16.95 (15.79, **26.22**) |
|  | SupFam | 6.97 | 5.47 | <u>23.73</u> | 19.31 | **25.80** | 21.76 | 15.85 (15.4, 11.87) | 17.52 (17.82, 20.38) |
| FlexNEQ | Fold | 5.71 | 12.98 | 35.95 | 16.41 | <u>45.08</u> | **49.64** | 15.41 (17.13, 17.37) | 18.68 (21.43, 18.95) |
|  | SupFam | 2.60 | 12.50 | 35.61 | 16.17 | <u>45.43</u> | **50.15** | 16.31 (18.33, 16.82) | 21.51 (24.22, 23.48) |
| Average Spearman's $\rho$% | | 7.80 | 9.84 | 33.49 | 21.81 | <u>37.35</u> | **38.08** | 16.61 (17.89, 21.36) | 20.84 (22.73, 23.89) |

Table 7 indicates that, under the PST-standard probe, our continuous embeddings capture non-trivial functional and physicochemical signals but remain behind sequence- and function-trained baselines on several functional-site tasks, particularly on harder SupFam splits. Notably, increasing probe expressiveness substantially narrows this gap: both the deeper MLP and the single-block transformer yield large gains, and the transformer probe produces the strongest overall results for both model sizes. This suggests that a significant portion of the task-relevant information is present in the learned latent space but is more readily extracted when the evaluation head can perform limited contextual aggregation across residues. Interestingly, despite weaker reconstruction fidelity, the Lite variant can match or slightly exceed the Large model under stronger probes, further supporting the view that reconstruction accuracy and downstream extractability are not strictly coupled.

For unsupervised conformational benchmarks (Table 8), our model achieves the best Apo–Holo correlations among the compared structure tokenizers, highlighting that the learned representations capture meaningful structural variation across conformational states. This suggests that our representation is particularly well-suited for applications demanding fine-grained structural discrimination rather than broad functional categorization. On fold-switching, our results remain competitive but trail the strongest baseline.

**Table 8:**
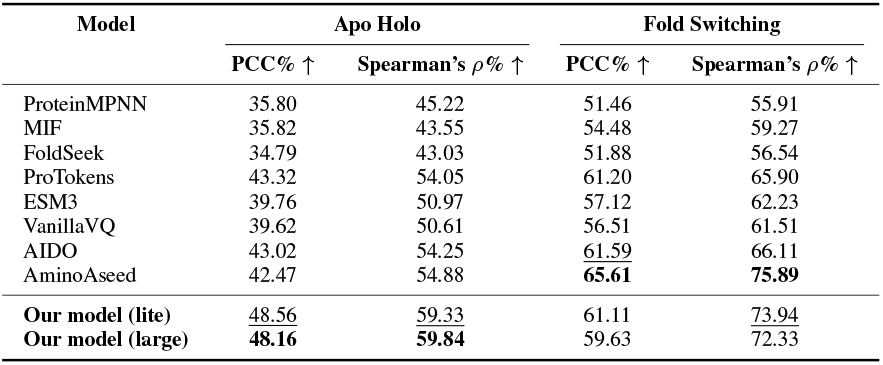
Performance on unsupervised Apo Holo and Fold Switching benchmarks.

**Table 9:** Training hyperparameters for Large (Stages 1 & 2) and Lite models.

| Hyperparameter | Large (Stage 1) | Large (Stage 2) | Lite |
| --- | --- | --- | --- |
| Optimizer type | AdamW | AdamW | AdamW |
| $\beta_1$ | 0.9 | 0.9 | 0.9 |
| $\beta_2$ | 0.98 | 0.98 | 0.98 |
| $\epsilon$ | $10^{-7}$ | $10^{-7}$ | $10^{-7}$ |
| Weight decay | $10^{-3}$ | $10^{-3}$ | $10^{-2}$ |
| Gradient clipping (max $L_2$ norm) | 1.0 | 1.0 | 1.0 |
| Gradient accumulation | 1 | 1 | 1 |
| 8-bit parameters | ✗ | ✓ | ✓ |
| Batch size (per GPU) | 16 | 4 | 16 |
| Learning-rate strategy | Cosine annealing | Cosine annealing | Cosine annealing |
| Base learning-rate | $1e-4$ | $5e-5$ | $1e-4$ |
| Min learning-rate | $1e-6$ | $1e-6$ | $1e-6$ |
| Warm-up steps | 16 000 | 16 000 | 4 096 |
| Total steps | 375 000 | 1 500 000 | 284 000 |
| MSE coefficient ($\lambda_{\text{MSE}}$) | $1 \times 10^{-3}$ | $5 \times 10^{-3}$ | $1 \times 10^{-3}$ |
| Backbone distance coefficient ($\lambda_{\text{dist}}$) | $10^{-2}$ | $10^{-2}$ | $10^{-2}$ |
| Backbone direction coefficient ($\lambda_{\text{dir}}$) | $5 \times 10^{-2}$ | $5 \times 10^{-2}$ | $5 \times 10^{-2}$ |

**Table 10:**
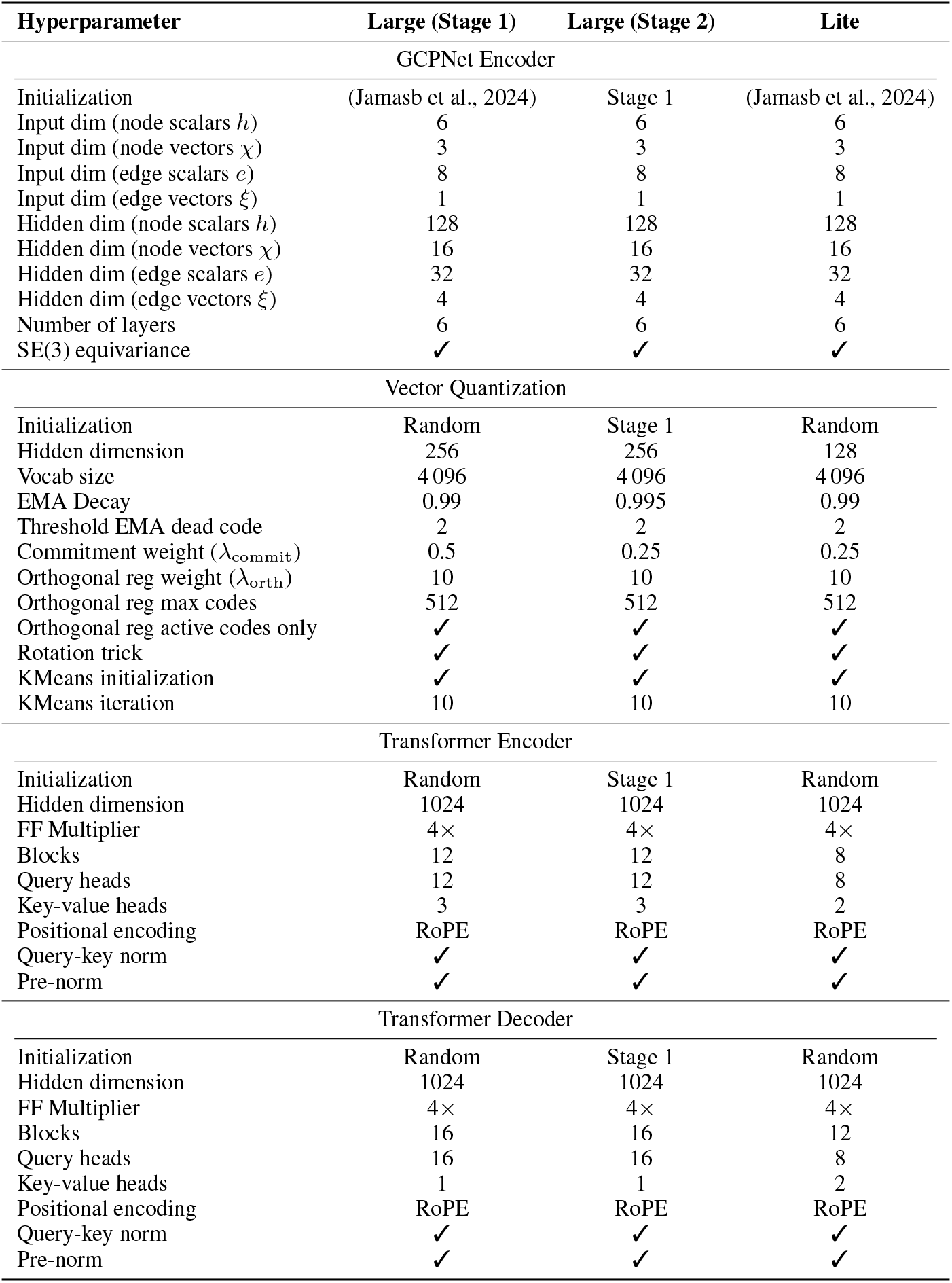
Configurations of GCP-VQVAE architectures (Large Stages 1 & 2, and Lite).

## 6 Discussion and Future Work

GCP-VQVAE delivers high-fidelity reconstruction across diverse suites of benchmarks (Figure 1A), with TM-scores typically ≥ 0.98 and mean RMSDs in the 0.40–0.80 Å range on CAMEO2024, CASP14/15/16, and similar performance on our internal Test A/B/C splits (e.g., 0.40–0.51 Å RMSD). Crucially, relative to the current method AIDO (Zhang et al., 2024), GCP-VQVAE not only yields lower reconstruction error but also reduces inference time by 408× for the Large model and 530× for the Lite variant, while remaining robust to structural noise (Appendix Figure 16).

On our diverse validation set, although overall reconstruction is strong, we observe a mild RMSD drift with sequence length (Figure 7). We attribute this to length imbalance in training; short chains dominate, leaving the model relatively underexposed to long-range constraints and rare structural motifs. Similarly, on the zero-shot set of 1 938 newly deposited experimental monomers, the model maintains strong generalization with mean RMSD 0.8193 Å and TM-score 0.9673. Our diagnostics indicate that degradation is driven primarily by the ratio and count of missing coordinates (Appendix Figure 10, 11), and secondarily by sequence length (Appendix Figure 9), similar to our interpretation of the validation results. Consistent with this, sequences without missing residues exhibit substantially lower errors than those with gaps (Table 5).

Comparing reconstruction and representation trends across tokenizers (Tables 2 and 7), we observe that high reconstruction accuracy does not necessarily imply superior downstream representation quality. For example, AminoAseed exhibits strong downstream performance yet very poor reconstruction, whereas our Large model achieves state-of-the-art reconstruction while the Lite model can yield stronger supervised transfer under richer probes. This highlights that reconstruction fidelity and downstream extractability are partially decoupled objectives, and that probe-based evaluation provides a complementary view of representation quality beyond reconstruction metrics alone.

Taken together, these results position GCP-VQVAE as a high-accuracy, fully open structure tokenizer that is both robust under distribution shift and practically usable at scale. Beyond reconstruction, our codebook analysis indicates that the learned tokens form a structured and highly utilized geometric vocabulary (Table 6), supporting the view that GCP-VQVAE learns a coherent “syntax” for protein backbones. Moreover, the Ramachandran analysis (Figure 3) suggests that individual tokens can span multiple local conformational regions rather than mapping to a single narrowly defined geometry, indicating that codes represent broader structural modes whose precise realization is resolved by context during decoding. We next highlight practical applications enabled by this geometry-complete discrete language.

Several avenues look promising to explore: scaling GCP-VQVAE tokenization to multi-chain complexes, enriching tokens with side-chain information, and evaluating GenAI-friendliness of this new language to unify sequence and structure generation via autoregressive protein language models (PLM).

### 6.1 Applications

For the proposed GCP-VQVAE architecture, we see the following applications:

#### Structure compression

Using a 4 096-entry codebook yields about ∼ 24 × compression of backbone coordinates for a 512-residue monomer (Appendix D). Given our low RMSDs of ∼ 0.4–0.8 Å (Table 2), this is a practical lossy backbone codec. Further gains are possible by (i) allowing a single code to represent short residue spans (e.g., 2–4 amino acids), reducing the token rate while the decoder expands span-tokens to per-residue poses, and/or (ii) increasing the codebook size (e.g., 8k–64k) to lower per-token quantization error so that fewer tokens are needed at a target distortion. These options trade bitrate against utilization and latency, and remain compatible with our VQ training approach.

#### Structure comparison

Our discrete geometry language converts backbones into high-resolution token strings that can be indexed (Altschul et al., 1990; Ma et al., 2002) and aligned with dynamic programming over a learned substitution matrix between codes. Compared to fixed 3Di alphabets as in FoldSeek, a learned 4 096-code vocabulary; optionally make it with side-chain–aware, captures finer local shape and orientation, enabling more accurate substructure and whole-chain comparisons.

#### Downstream 3D learning representation

De-quantized code embeddings yield continuous, structure-aware features that can serve as a compact intermediate representation for a range of downstream 3D learning tasks (Yuan et al., 2025b). In practice, these embeddings can be used as fixed inputs for supervised residue-level prediction (e.g., functional site annotation), physicochemical property regression, and unsupervised conformational analysis, providing a reusable structural featurization that bridges discrete token sequences and continuous geometric learning. This interface enables transferring geometric information from a high-fidelity tokenizer into lightweight task-specific heads, and supports applications such as structure-aware retrieval, clustering, and structure-conditioned modeling without operating directly on raw coordinates.

#### Generative modeling

Because our structure representation is discrete and pose-invariant, it slots directly into autoregressive PLM pre-training: tokens can be interleaved with amino-acid symbols, letting us exploit next-token scaling laws observed in autoregressive Large Language Models (LLM) (Kaplan et al., 2020). This unifies sequence and structure in a single model, enabling controllable generation via simple conditioning; e.g., sequence-given-structure and structure-given-sequence modes. Structure tokens serve as an explicit geometric prior for PLMs (Hayes et al., 2025; Su et al., 2023), potentially enabling more control over protein design. Moreover, the same machinery supports higher-throughput structure prediction from sequence by first predicting structural tokens and then mapping them to backbones with GCP-VQVAE’s decoder (Pourmirzaei et al., 2025; Lu et al., 2024; Chen et al., 2024).

### 6.2 Limitations

#### Backbone-only discretization

Our tokenizer operates solely on backbone coordinates (N–C_*α*_– C) and does not incorporate side-chain geometry into the discrete vocabulary. While this design enables stable training and high-fidelity backbone reconstruction, it omits chemically informative details such as rotamer states, functional group orientation, and local packing interactions that are often critical for binding and catalysis.

#### Reconstruction–representation gap

Although GCP-VQVAE achieves strong reconstruction accuracy, representation transfer on supervised functional benchmarks is comparatively lower, indicating that reconstruction fidelity alone does not ensure that downstream-relevant information is easily extractable from the learned embeddings. Notably, this gap narrows substantially when using more expressive yet lightweight probes (e.g., a deeper MLP or a single-block transformer), indicating that a meaningful fraction of task-relevant signal is present but may require limited contextual aggregation to decode. Moreover, the current training objective is purely reconstruction-driven and does not incorporate auxiliary supervision (e.g., inverse folding, distillation from pretrained protein encoders, or other multi-task signals) that could enrich the semantic content of the vector-quantized embedding space.

#### Monomer-only scope

We train and evaluate GCP-VQVAE exclusively on single-chain proteins and do not explicitly support multi-chain tokenization. As a result, the learned discrete language does not capture inter-chain geometry or interface-driven constraints that arise in complexes, limiting direct applicability to settings where quaternary structure plays a central functional role.

## 7 Conclusion

In this work, we have introduced GCP-VQVAE, a state-of-the-art open-source protein structure tokenizer. GCP-VQVAE offers improved structure reconstruction quality and generalization compared to prior methods, which could enable new (sequence and structure-based) predictive and generative modeling applications in future work. In the end, we released the GCP-VQVAE source code, pretrained checkpoints, curated datasets, and data preprocessing pipeline to the research community.

## Acknowledgments

This research used NVIDIA GPU resources awarded via the NAIRR Pilot (Allocation: NAIRR240246). We thank NAIRR and NVIDIA for supporting the computations underlying our results.

## Appendix

### A Methods

The proposed architecture leverages two main parts: (1) a GCPNet encoder to encode backbone coordinates into embeddings, and (2) a transformer-based VQVAE, which discretizes backbone embeddings and then converts them back into 3D coordinates.

#### A.1 gcpnet Encoder

GCPNet (Morehead & Cheng, 2024b) extends the scalar–vector message-passing philosophy of GVP-GNN to a geometry-complete, SE(3)-equivariant encoder. Every atom *i* in a molecular graph *G* = (*V, E*) carries scalars 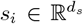 and row-wise vectors 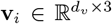 that ro tate as 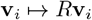 under *g* = (*R*, **t**) ∈ SE(3), while each edge (*i, j*) stores analogous features *s*_*ij*_, **v**_*ij*_ and the relative displacement **r**_*ij*_ = **x**_*j*_ − **x**_*i*_. Before any update, the model attaches to every edge a right-handed orthonormal frame *F*_*ij*_ = [**a**_*ij*_, **b**_*ij*_, **c**_*ij*_] ∈ SO(3) with **a**_*ij*_ = **r**_*ij*_*/* ∥ **r**_*ij*_ ∥; this frame supplies a reference for chirality and orientation that GVP-GNN lacks.

The core computation is a Geometry-Complete Perceptron (GCP) micro-step that first down-scales the vectors and then projects them into the local frame to extract nine orientation-aware features. Denoting **z**_*ij*_ = *W*_*d*_**v**_*ij*_ and vec() the row-wise vectorization, the joint update mixes the old scalars with their orientation signatures (the frame-projected vectors and their norms) and gates the vectors through a learnable row-wise gate to preserve equivariance; *ϕ*_*s*_ is an MLP and *σ* denotes a learnable gating function that need not be a fixed sigmoid (Equation 3).

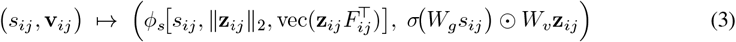

In the node update, the orientation features are averaged over neighbors. A sequence of such micro-steps, wrapped by a residual shortcut, forms a GCPConv edge block. After aggregating messages *m*_*i*_ = ∑_*j* ∈ 𝒩 (*i*)_ GCPConv(*i, j*), a gated scalar–vector MLP updates the node features, and stacking *L* layers yields an invariant backbone. When tasks require coordinates, each layer appends an equivariant displacement head whose output is a learned 3-dimensional vector; this vector is added residually to **x**_*i*_ and re-centered to ensure translation invariance, enabling force or trajectory prediction without breaking equivariance.

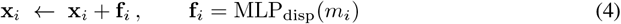

Equations 3 and 4 commute with every rigid motion, making the encoder strictly SE(3)-equivariant, while projection through *F*_*ij*_ preserves the complete set of edge orientations so that the latent remains *geometry-complete* at any depth. Eliminating the frame (*F*_*ij*_ = *I*), replacing the orientation features by ∥ **v**_*ij*_ ∥ _2_, and dropping the coordinate head restores a frame-free, E(3)-equivariant network akin to GVP-GNN—thereby isolating the contributions responsible for our empirical gains. Because GCPNet keeps directional and chiral cues that GVP-GNN discards, it supports optional equivariant coordinate updates for force or dynamics prediction and attains improved performance across invariant (e.g., binding affinity; Morehead & Cheng (2024b)), equivariant (e.g., force regression; Morehead & Cheng (2024b)), and coordinate-generative (e.g., molecular diffusion; Morehead & Cheng (2024a)) tasks with only modest extra computation.

#### A.2 VQVAE

The transformer–based VQVAE employed in this work is organized into the classical three-stage pipeline of *encoder, vector-quantization*, and *decoder*. The encoder processes the given embeddings into a sequence of latent vectors, the quantizer discretizes these latents, and the decoder reconstructs the original signal from the resulting code indices.

Both stacks adopt a lightweight pre-layer-normalized transformer that integrates several recent efficiency upgrades: (i) Pre-LayerNorm places the LayerNorm before each sub-block (Xiong et al., 2020), which keeps activations in a well-behaved range throughout the network, reduces gradient-scale drift, and therefore allows training with larger learning rates and much milder warm-up schedules; (ii) separately normalizes query and key vectors before computing attention logits (Henry et al., 2020). This prevents overly large dot-products, stabilizes attention distributions, and mitigates soft-max saturation, especially in low-resource or small-batch training scenarios; (iii) Grouped-Query Attention shares key–value projections across groups of query heads (Ainslie et al., 2023), reducing both memory and compute without harming quality; (iv) Rotary Positional Embeddings (RoPE) inject relative-position information by applying position-dependent planar rotations to each query–key pair (Su et al., 2024), letting the model generalize to much longer sequences with virtually no extra computational cost; (v) We remove bias terms from projection and feed-forward layers, an established simplification that has negligible effect on accuracy while trimming a fraction of parameters and FLOPs.

In the architecture, the vector quantization layer provides the discrete bottleneck between the afore-mentioned encoder and decoder. While fixed quantizers such as finite scalar quantization (FSQ) (Mentzer et al., 2023) can simplify optimization and offer strong rate–distortion behavior, our target application emphasizes a discrete *representation* that can serve as a reusable interface for both reconstruction and downstream generative modeling. Learned VQ codebooks provide an explicit set of trainable embeddings whose geometry and usage statistics can be shaped by objectives beyond pixel/coordinate reconstruction, which is important when tokens are later modeled by autoregressive or masked-token transformers.

This design choice is supported by recent findings in visual tokenization: tokenizers that improve *semantic* fidelity (e.g., by distilling from strong understanding encoders) yield better generation quality, indicating that learned discrete spaces can capture information that is directly useful for generative modeling (Wang et al., 2024a). Furthermore, work on the compression–generation trade-off shows that stronger generation can emerge from token spaces that are intentionally easier for a stage-2 model to learn, motivating tokenizer training recipes that go beyond pure reconstruction (Ramanujan et al., 2024). Recent studies demonstrated that the VQ-based tokenizers are scalable and can reach full (100%) codebook utilization and avoid classic collapse/under-use pathologies (Chang et al., 2025). Given that our training achieves near-complete codebook utilization on validation, we retain learnable VQ rather than switching to a fixed quantizer, preserving the ability to incorporate auxiliary semantic constraints into the token space when needed.

Before training starts, we run *k*-means on the encoder outputs of the first mini-batch to seed the codebook as displayed in Equation 5, where *Z*^(0)^ ⊂ ℝ^*d*^ are the features, *K* the codebook size, and *T* the number of Lloyd iterations. Empirically, this step mitigates early code collapse and improves utilization when *K* is large.

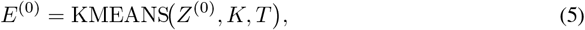

Given an encoder vector **z** ∈ ℝ^*d*^, quantization proceeds by nearest-neighbor lookup, Equation 6, which is identical to the vanilla VQ rule.

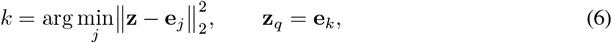

To transmit gradients through this non-differentiable operation, instead of using the straight-through estimate (STE) introduced in Van Den Oord et al. (2017), we adopt the rotation trick (Fifty et al., 2024). During back-propagation, the Jacobian is replaced by Equation 7, where *R* is the shortest-arc rotation aligning the unit vectors 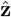 and 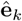 This modification embeds both angular and magnitude mismatch into the back-propagated signal, yielding faster convergence and richer code usage in practice.

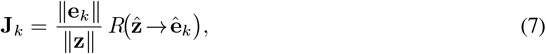

The codebook updates and the encoder commitment follow the standard VQ losses (Equation 8), with *β* balancing the commitment pressure.

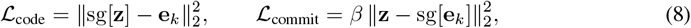

To encourage diverse and well-separated embeddings, we regularize the codebook with the Frobenius norm (Equation 9) weighted by *λ*_orth_. This orthogonality constraint spreads codes over the hypersphere and curbs under-utilization, which is a common failure mode in large books.

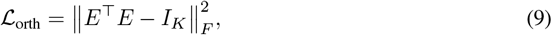

To translate the decoder’s abstract embeddings into a physically meaningful 3D structure, we utilize a 6D rotation head on top of the decoder. This module’s primary purpose is to provide a stable and continuous parameterization of 3D rotations and translations, which is crucial for effective training of deep neural networks. This approach, notably used in AlphaFold2 and supported by the findings of Zhou et al. (2019) on rotation representations, avoids the well-known issues of Gimbal lock in Euler angles and the double-cover ambiguity of quaternions.

Intuitively, the head operates by predicting an intermediate representation for each residue, comprising two 3D direction vectors and a translation. The direction vectors are deterministically converted into a stable rotation matrix via the Gram-Schmidt process, while the translation is scaled by a hyper-parameter, *α*, to an arbitrary range (e.g., Å). This resulting rigid transformation acts as an update, which is composed with the residue’s running pose. The final backbone coordinates are then generated by applying this new, refined pose to a fixed local atomic template (**X**^local^). This iterative process ensures the structure is built in a geometrically consistent and equivariant manner, with the full operational details provided in Algorithm 1.

The decoder is supervised with a weighted sum of three geometric objectives. Building on the loss terms defined in Algorithm 2 (distance, direction and aligned MSE) and the Kabsch alignment returned by Algorithm 3, we supervise the decoder with the weighted sum of Equation 10 as the reconstruction part (*L*_rec_) of the final loss. We emphasize that Kabsch alignment removes only global pose (proper rigid motion with det(*R*) = +1), while the distance and direction losses are defined on internal geometric relations and remain invariant to any global SE(3) transform.

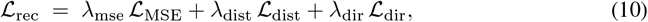

Here, ℒ _MSE_ is the mean-squared error between predicted and native backbones after Kabsch alignment, ℒ _dist_ penalises deviations in the 3*L* × 3*L* backbone distance matrix (clamped at 5 Å^2^), and ℒ _dir_ measures squared differences of pairwise dot-product tensors over the six orientation vectors per residue (clamped at 20). Finally, the overall objective optimized for each iteration is demonstrated in Equation 11 where *L*_rec_ is the decoder reconstruction.

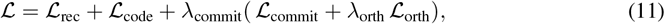

Algorithms 1–3 define the decoder head, geometric losses, and alignment used in training. Given per-residue embeddings, the 6D head projects to two direction vectors and a translation, constructs a proper rotation via Gram–Schmidt, composes this rigid update with the running pose, and applies it to a fixed (N–C_*α*_–C) template to produce backbone coordinates. Reconstruction is supervised by the weighted sum in Equation 10: (i) aligned MSE after optimal Kabsch alignment, (ii) a clamped backbone distance-matrix loss, and (iii) a clamped pairwise orientation (direction) loss built from six backbone vectors per residue.

##### Algorithm 1 Pseudocode for 6D rotation-based structure prediction

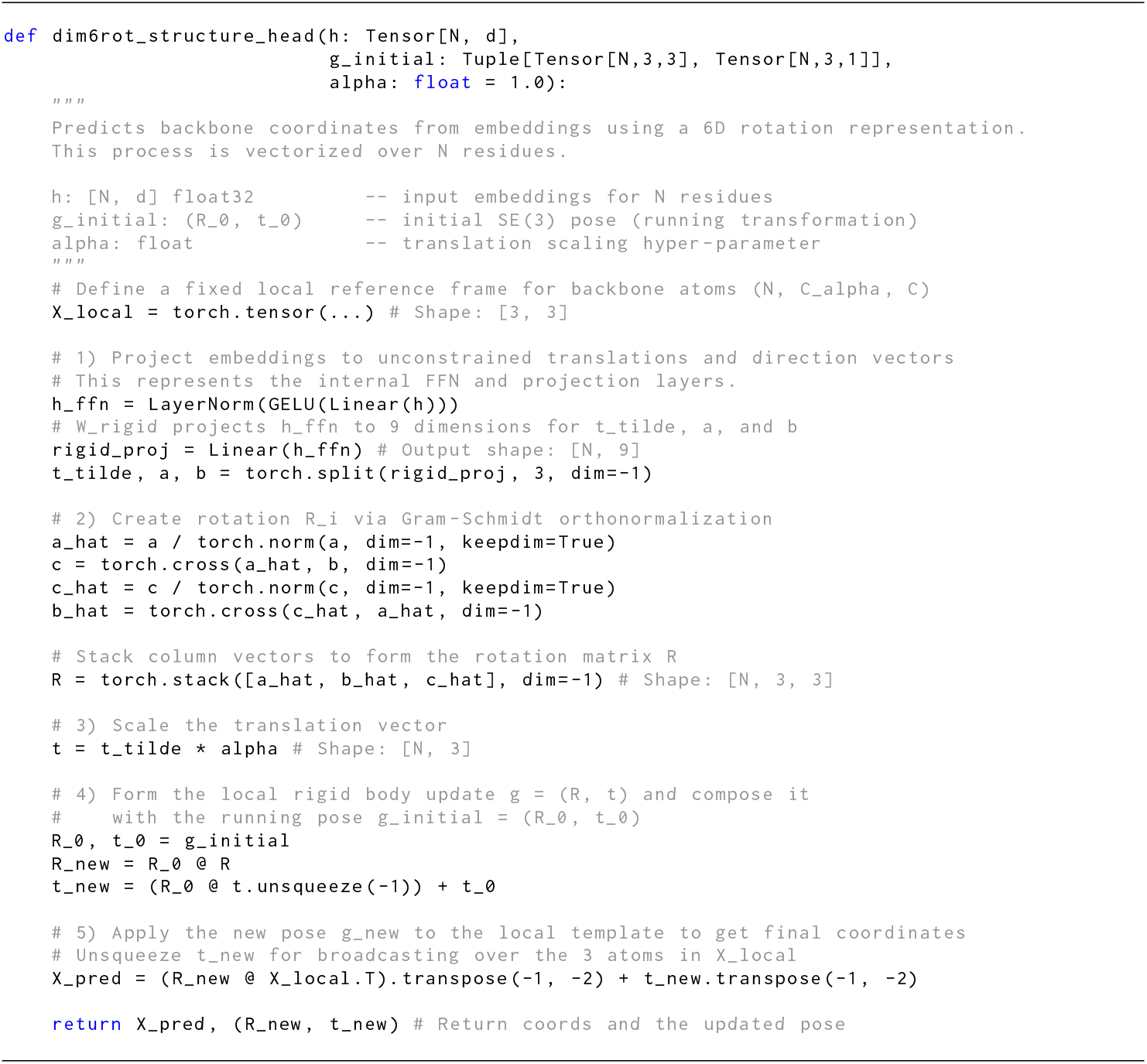

##### Algorithm 2 Pseudocode for backbone *distance, direction* and *MSE* losses

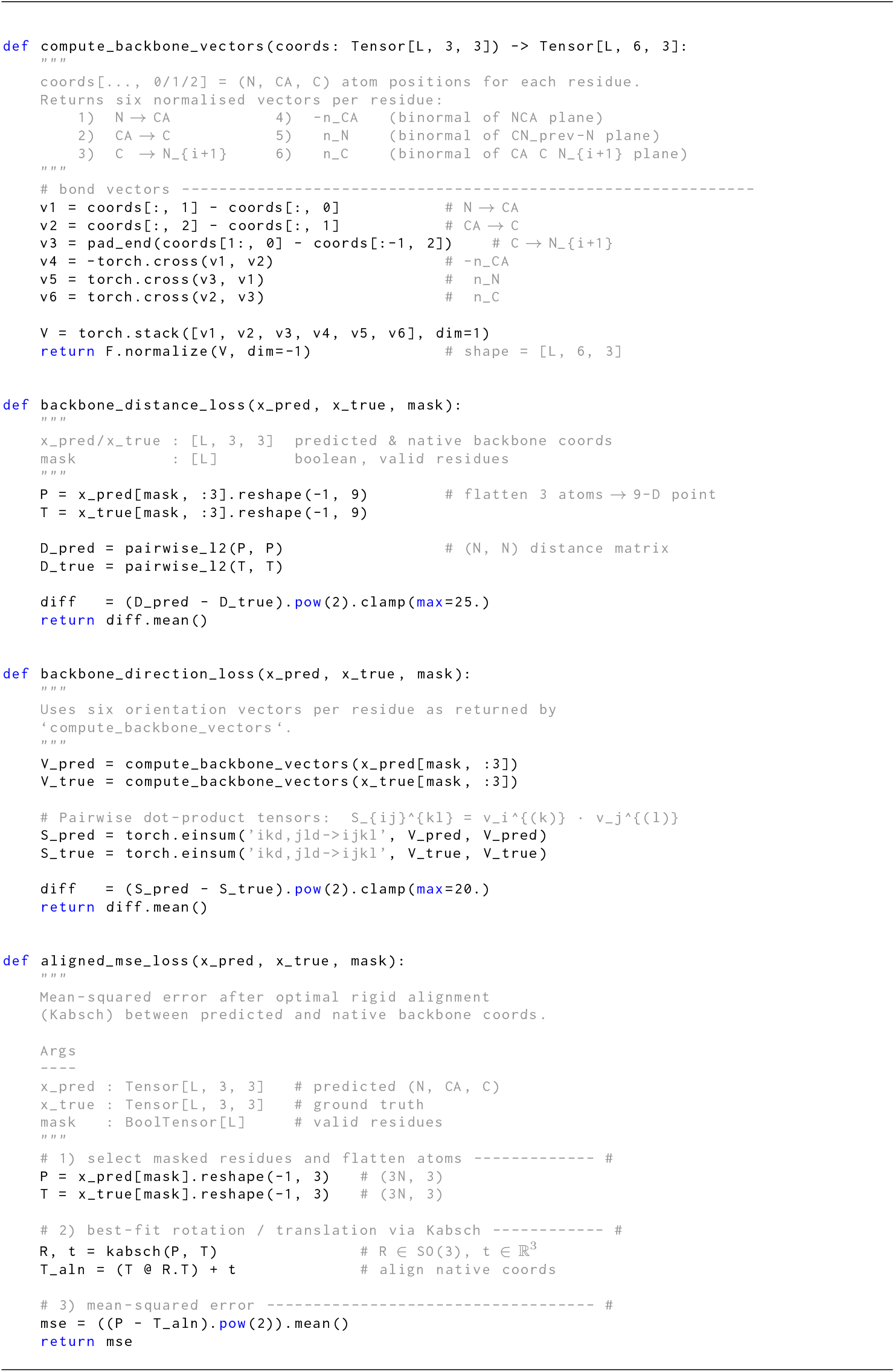

##### Algorithm 3 Pseudocode for rigid alignment via Kabsch

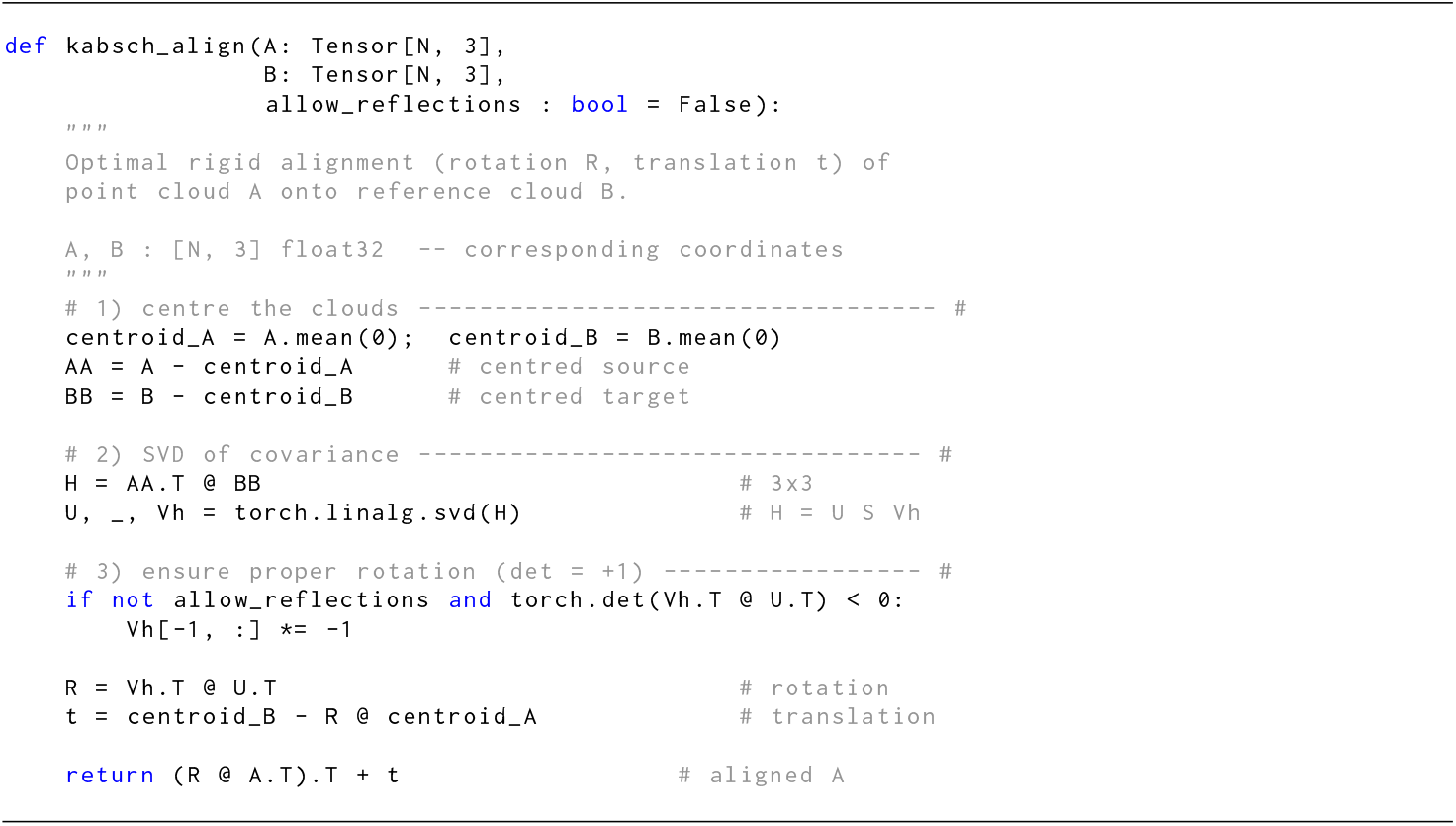

### B Datasets

*Test A (low-confidence)* evaluates robustness under structural uncertainty using a curated subset of AlphaFold2 (AF2) models for SwissProt proteins. We enumerated all AF2-predicted structures available for SwissProt entries in AFDB and retained only those with average pLDDT ≤ 70 (AFDB’s 0–100 scale), discarding higher-confidence models. The surviving records were *cross-referenced* against the PDB to annotate the presence of experimental structures for the same proteins. Multimer entries were decomposed into per-chain monomer samples, after which we removed chains shorter than 25 amino acids and retained only chains with fewer than 25 consecutive missing residues. A final deduplication step produced 33 112 single-chain samples for Test A.

*Test B (species-diverse, taxonomic shift)* assesses generalization under distributional shift in species rather than structure confidence. We assembled a broad species roster (expanded to 100 and finalized at 97 species) and targeted roughly 2 500 proteins per species from AFDB without pLDDT filtering to reflect natural variability, dropping species with fewer than 1 000 available structures. After aggregation, per-chain conversion, and deduplication, this suite comprised 19 353 single-chain samples.

*Test C (high-confidence, under-represented taxa)* measures performance on high-confidence structures from taxa under-represented in SwissProt. We selected species with low SwissProt coverage and required high average pLDDT (*>* 95). The selected taxonomies were cross-referenced against the PDB to find the presence of experimental structures for the proteins from those species. Applying the same preprocessing and deduplication as Test A above yielded 33 867 single-chain samples.

To obtain a balanced validation set independent of training, we first constructed three targeted hold-out suites and then set aside a fixed 2 000 randomly sampled chains from each to form a combined 6 000-chain validation set; the remaining examples constitute Test sets A, B, and C, respectively. The distribution of length sequences in the test sets is displayed in Figure 4.

**Figure 4.**
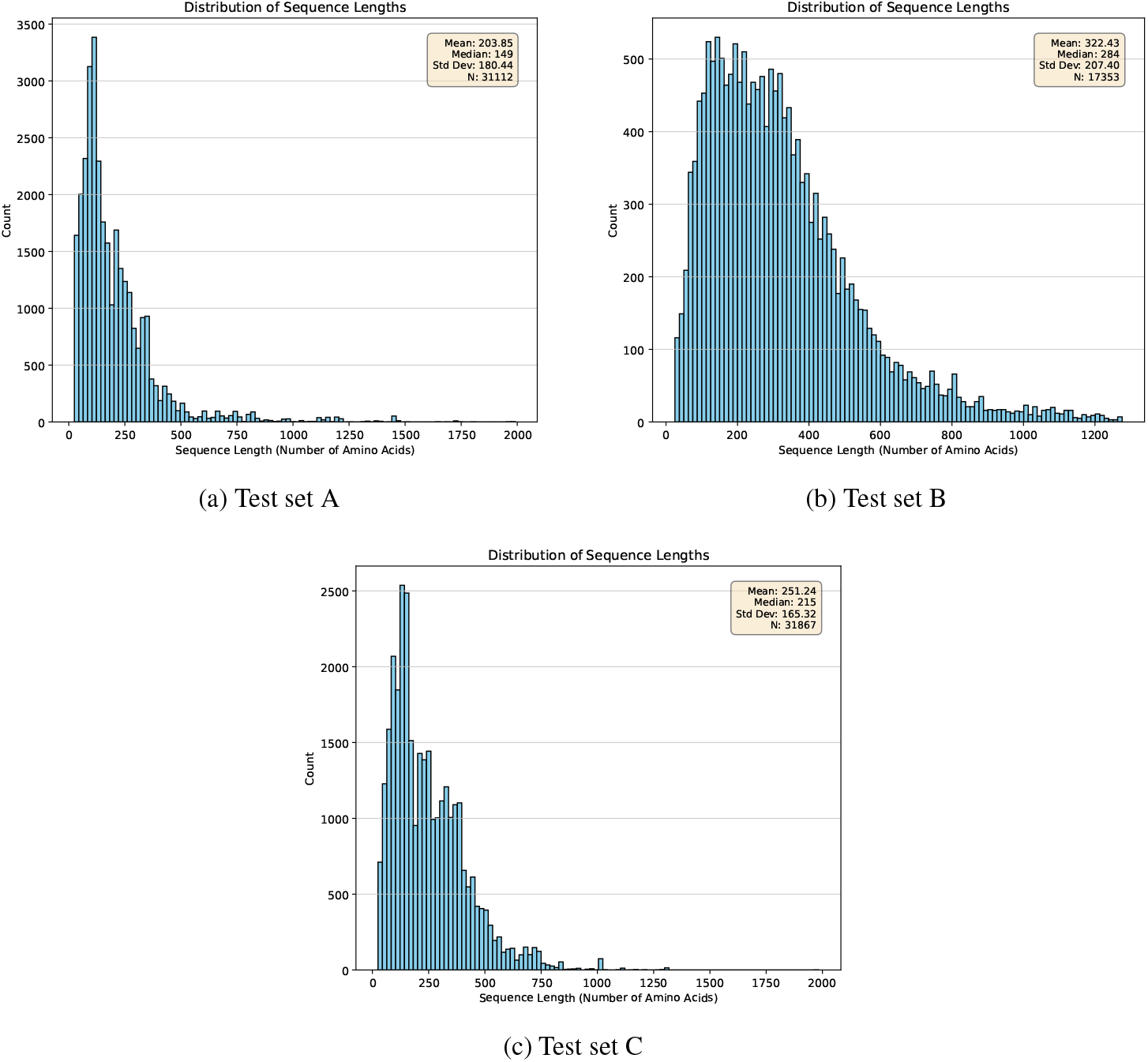
Distribution of sequence lengths in the three test sets. The majority of samples have sequence length *<* 1280 amino acids.

**Figure 5.**
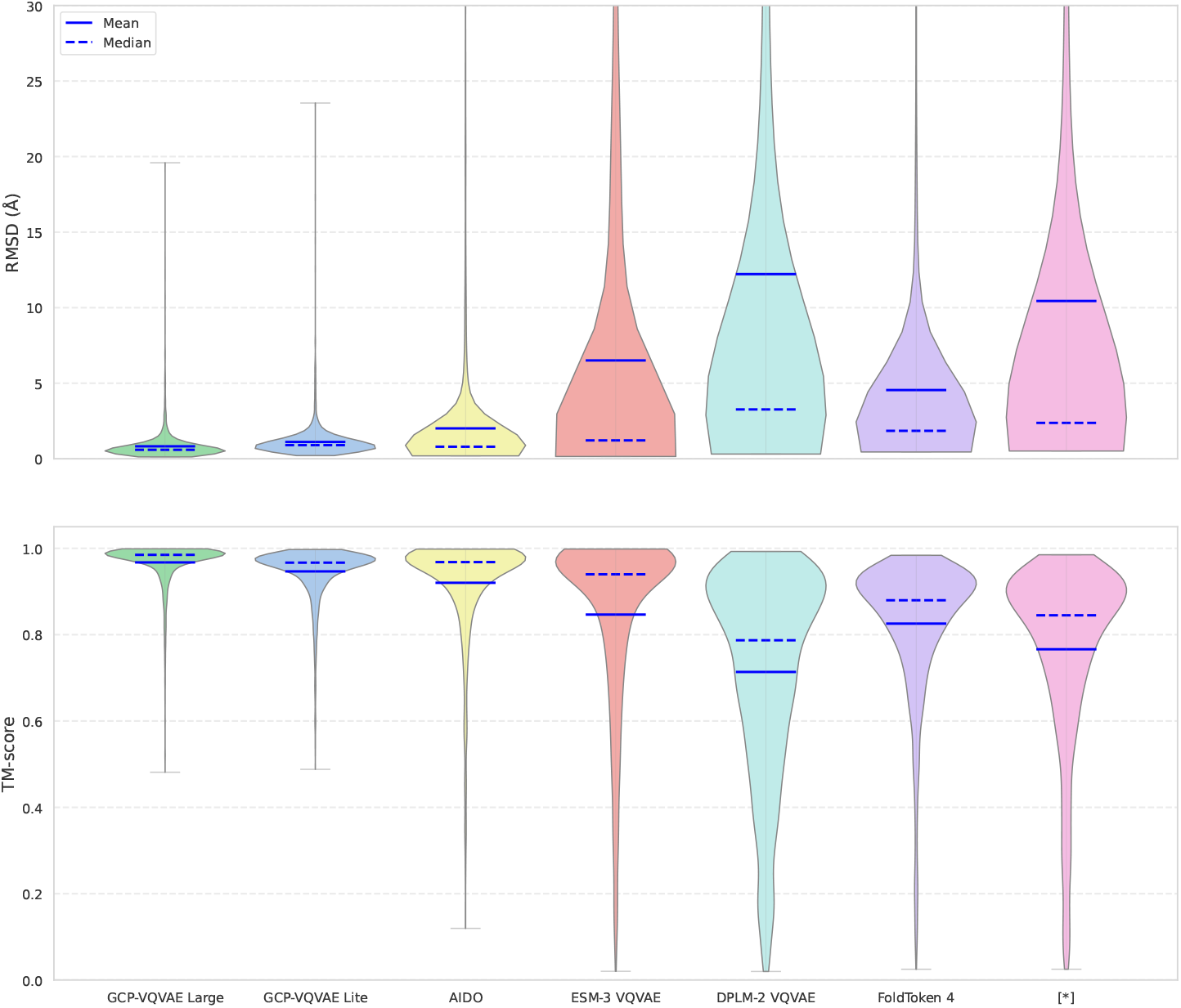
RMSD (Å) and TM-score error distributions on the zero-shot dataset. Gaujac et al. (2024) is shown as [∗]. For Gaujac et al. (2024), samples outside its supported length range (50–512 residues) are excluded; other methods use the full sets.

**Figure 6.**
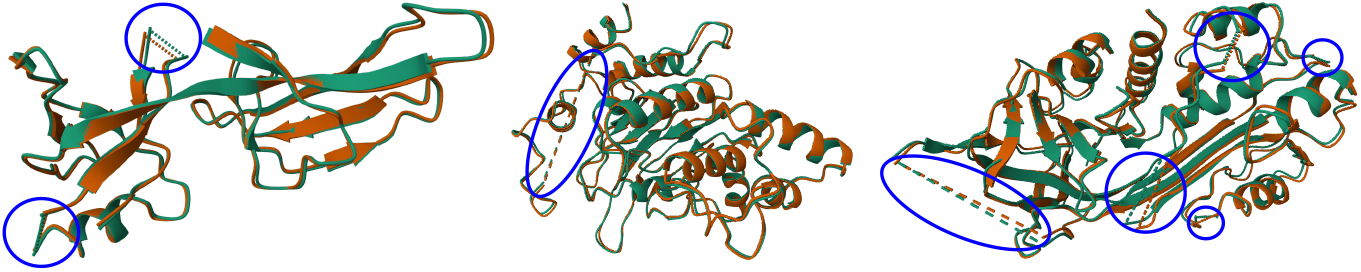
Superposition of GCP-VQVAE reconstructions (orange) with native backbones (green) for three proteins drawn from our external benchmark suites. Blue circles mark contiguous missing-residue segments (dashed guides span regions without deposited atoms); the model tracks the native structure outside the gaps.

**Figure 7.**
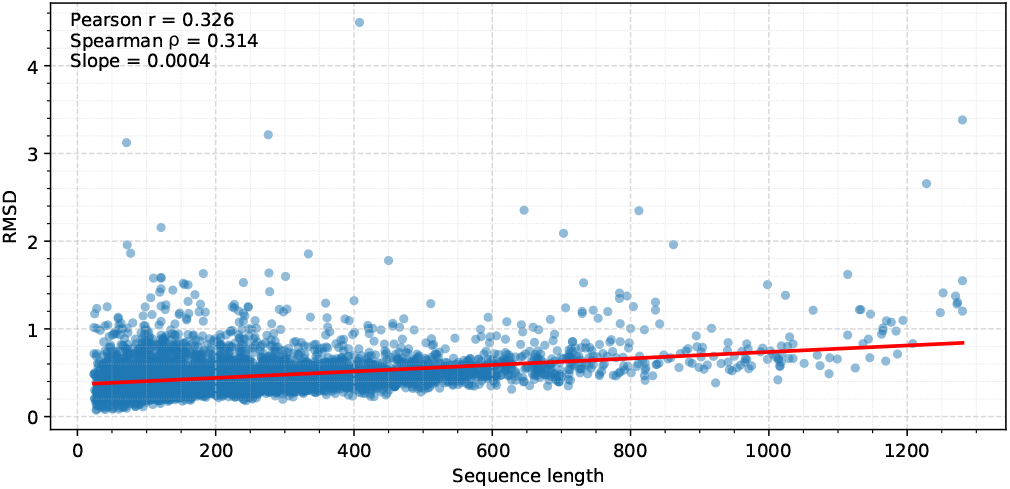
Sequence length vs. reconstruction error (validation set). Each point is one protein; the red line is a least-squares fit. The dependence on length is modest (Pearson *r* = 0.326, Spearman *ρ* = 0.314) with a small slope of ≈ 4 × 10^−4^ Å/residue (∼ 0.4 Å per 1k residues).

From the zero-shot set, we defined interpretable structural subsets based on each protein’s secondary-structure composition. Coil-rich (D) includes sequences with at least 60% coil/disordered content. Mainly *α* (A) requires an *α*-helical fraction of at least 45% and an *α* − *β* difference of at least 15 percentage points; mainly *β* (B) requires a *β*-strand fraction of at least 25% and a *β* − *α* difference of at least 10 points. Balanced (AB) captures mixed folds with *α* ≥ 25%, *β* ≥ 15%, and |*α* – *β*| ≤ 10 points. We also report two data-quality subsets: sequences with no missing residues and sequences with at least one missing residue. Groups are defined independently (i.e., they may overlap).

### C Experiments

This subsection compiles all experimental assets: Table 9 lists the full optimization schedule and training hyperparameters for the Lite and Stages 1–2 of the Large model; Table 10 specifies the exact model configuration (GCPNet, VQ, and transformer stacks). Figure 5 reports RMSD error distributions on the zero-shot dataset across methods; Figure 6 shows qualitative robustness to contiguous missing-residue segments from benchmark structures; Figure 7 plots sequence length versus reconstruction RMSD with a least-squares fit on the validation set; Figure 8 visualizes the highest-RMSD (worst-case) examples per benchmark suite. Metrics use backbone atoms (C_*α*_) after Kabsch alignment, and statistics are over the full, unfiltered test sets.

**Figure 8.**
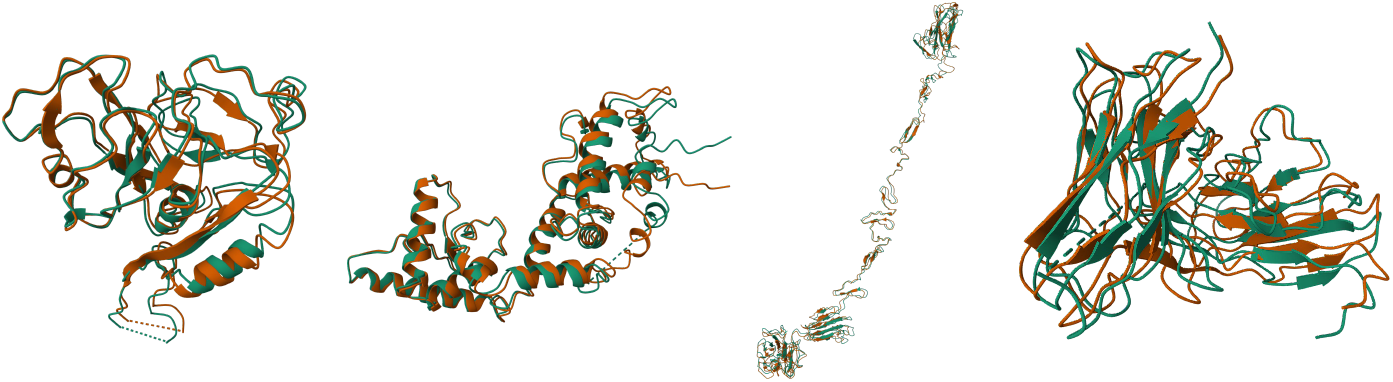
Superposition of GCP-VQVAE (orange) and native backbones (green) for the highest–backbone-RMSD example in each suite, left→right: CASP14, CASP15, CASP16, CAMEO2024.

**Figure 9.**
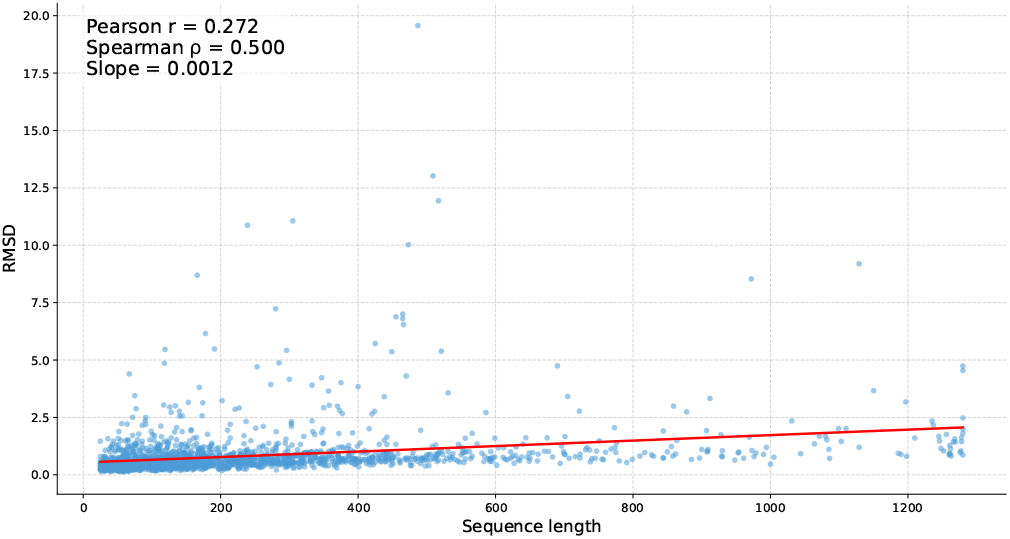
RMSD as a function of sequence length (number of amino acids) on the zero-shot set. Each dot is one protein. The red line is an ordinary-least-squares (OLS) fit; the panel reports Pearson *r*, Spearman *ρ*, and the fitted slope (RMSD units per residue).

**Figure 10.**
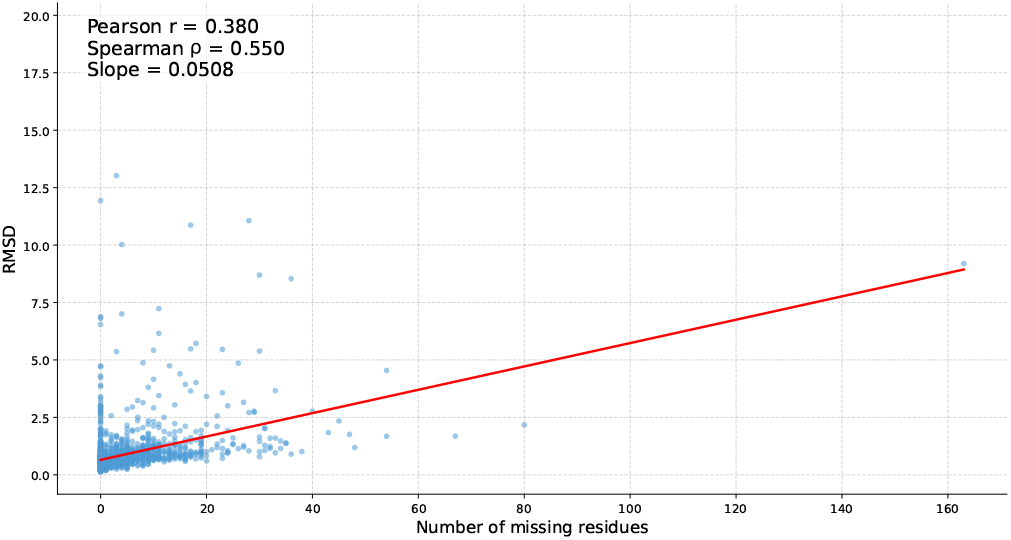
RMSD as a function of the number of missing residues on the zero-shot set. Points show individual proteins; the red line is an OLS fit. The panel reports Pearson *r*, Spearman *ρ*, and the fitted slope (RMSD units per missing residue).

**Figure 11.**
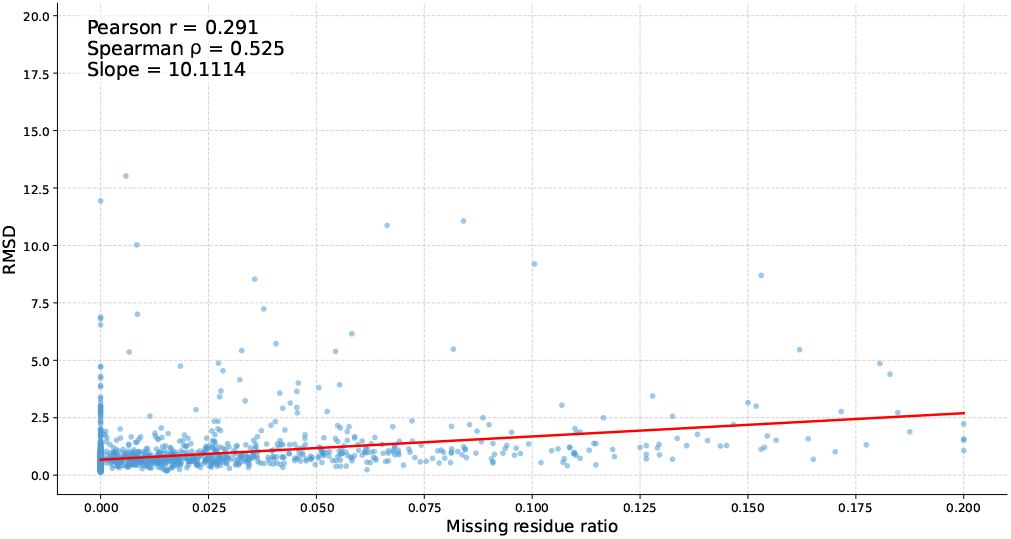
RMSD as a function of the missing-residue ratio on the zero-shot set. The red line is an OLS fit; the panel reports Pearson *r*, Spearman *ρ*, and the fitted slope (RMSD units per unit fraction; 1.0=100%).

**Figure 12.**
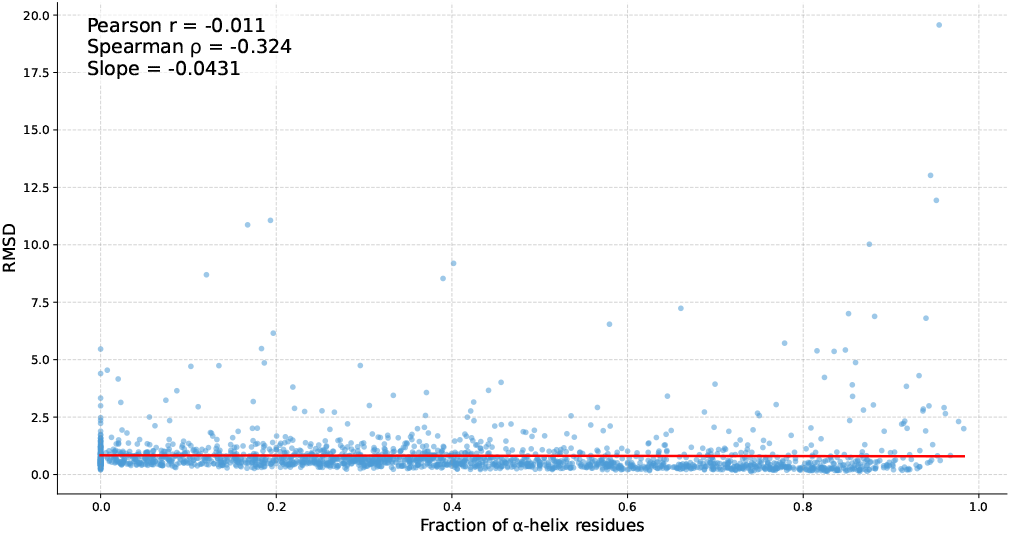
RMSD as a function of the fraction of *α*-helix residues on the zero-shot set. The red line indicates an OLS fit. The panel reports Pearson *r*, Spearman *ρ*, and the fitted slope (RMSD units per unit fraction; 1.0=100%).

**Figure 13.**
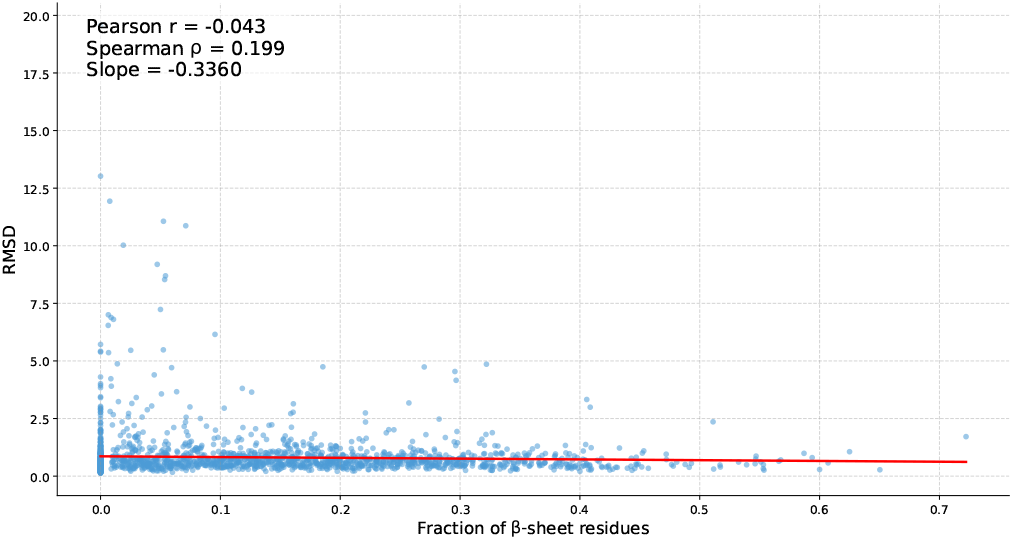
RMSD as a function of the fraction of *β*-sheet residues on the zero-shot set. The red line indicates an OLS fit. The panel reports Pearson *r*, Spearman *ρ*, and the fitted slope (RMSD units per unit fraction; 1.0=100%).

**Figure 14.**
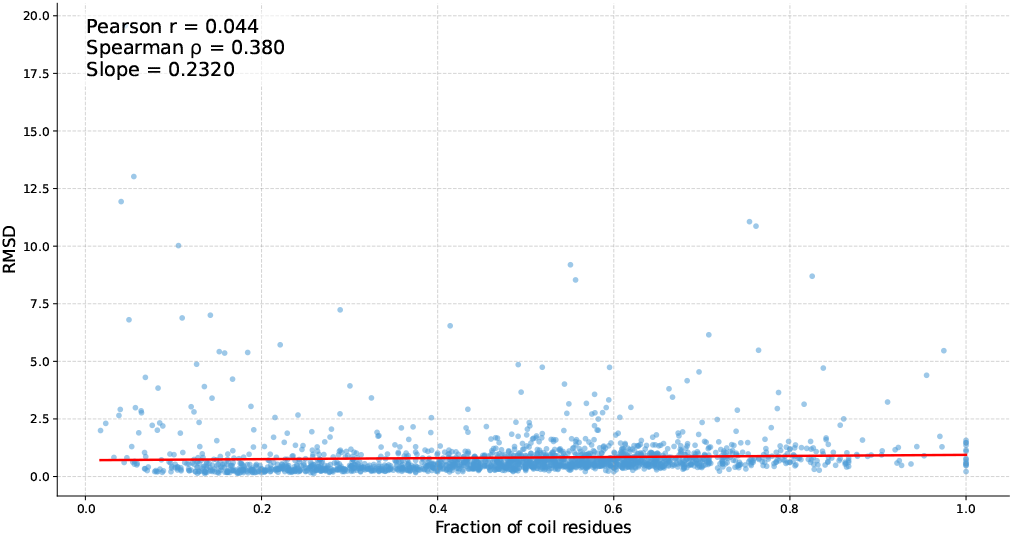
RMSD as a function of the fraction of coil residues on the zero-shot set. The red line indicates an OLS fit. The panel reports Pearson *r*, Spearman *ρ*, and the fitted slope (RMSD units per unit fraction; 1.0=100%).

**Figure 15.**
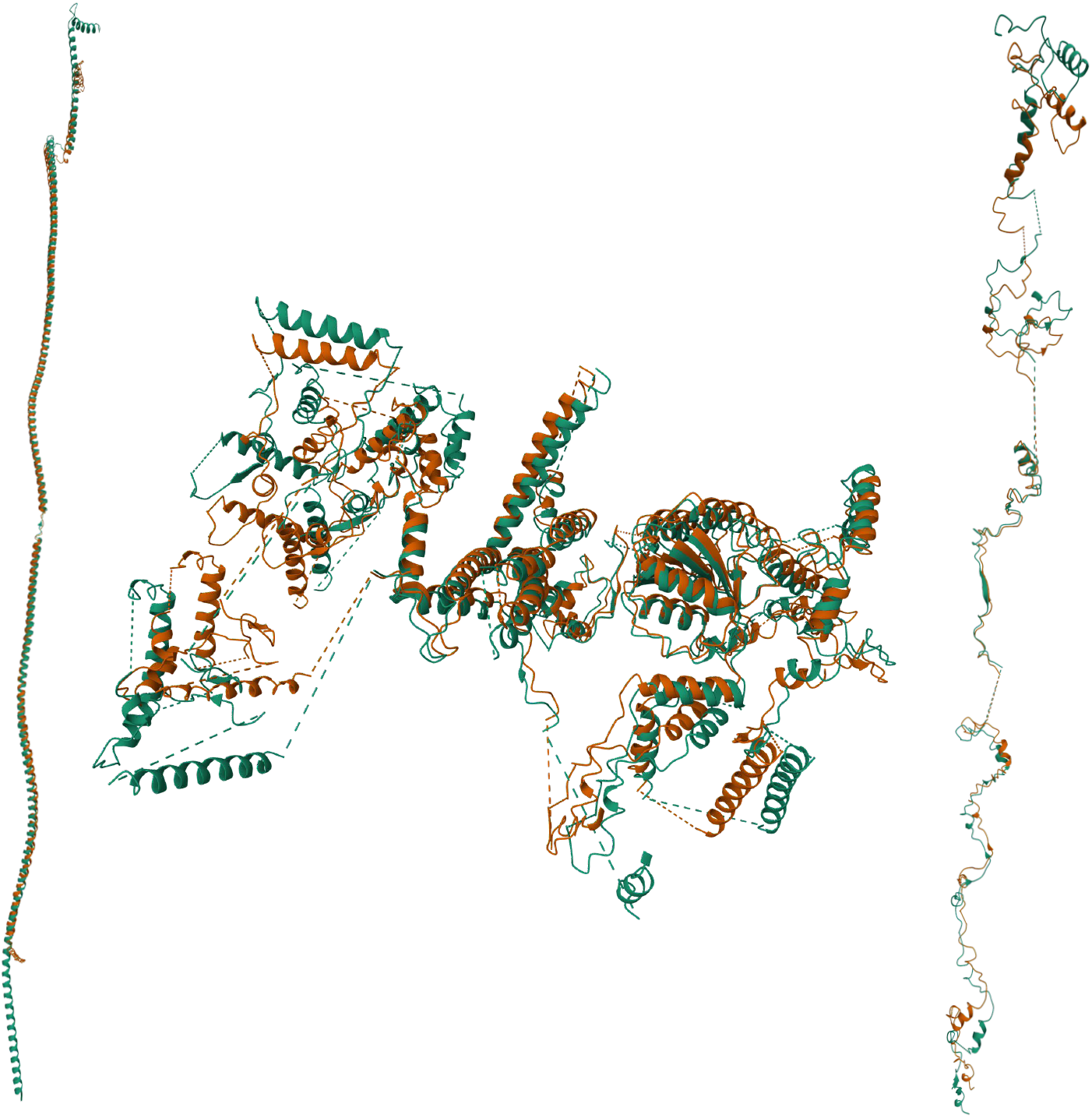
Worst zero-shot reconstructions (orange: GCP-VQVAE Large; green: native). Left: RMSD 19.568, TM-score 0.691, length 487, NaN residues 0. Middle: RMSD 9.1924, TM-score 0.7442, length 1129, NaN residues 163. Right: RMSD 11.0627, TM-score 0.4755, length 305, NaN residues 28. The middle and right cases exhibit an extreme number of NaN residues introduced by our current NaN-handling/augmentation pipeline, which forces the model to contend with long masked segments and degrades reconstruction accuracy; improving the NaN strategy (e.g., capping gap ratio, better segmentation, and length-aware masking) should mitigate this failure mode.

**Figure 16.**
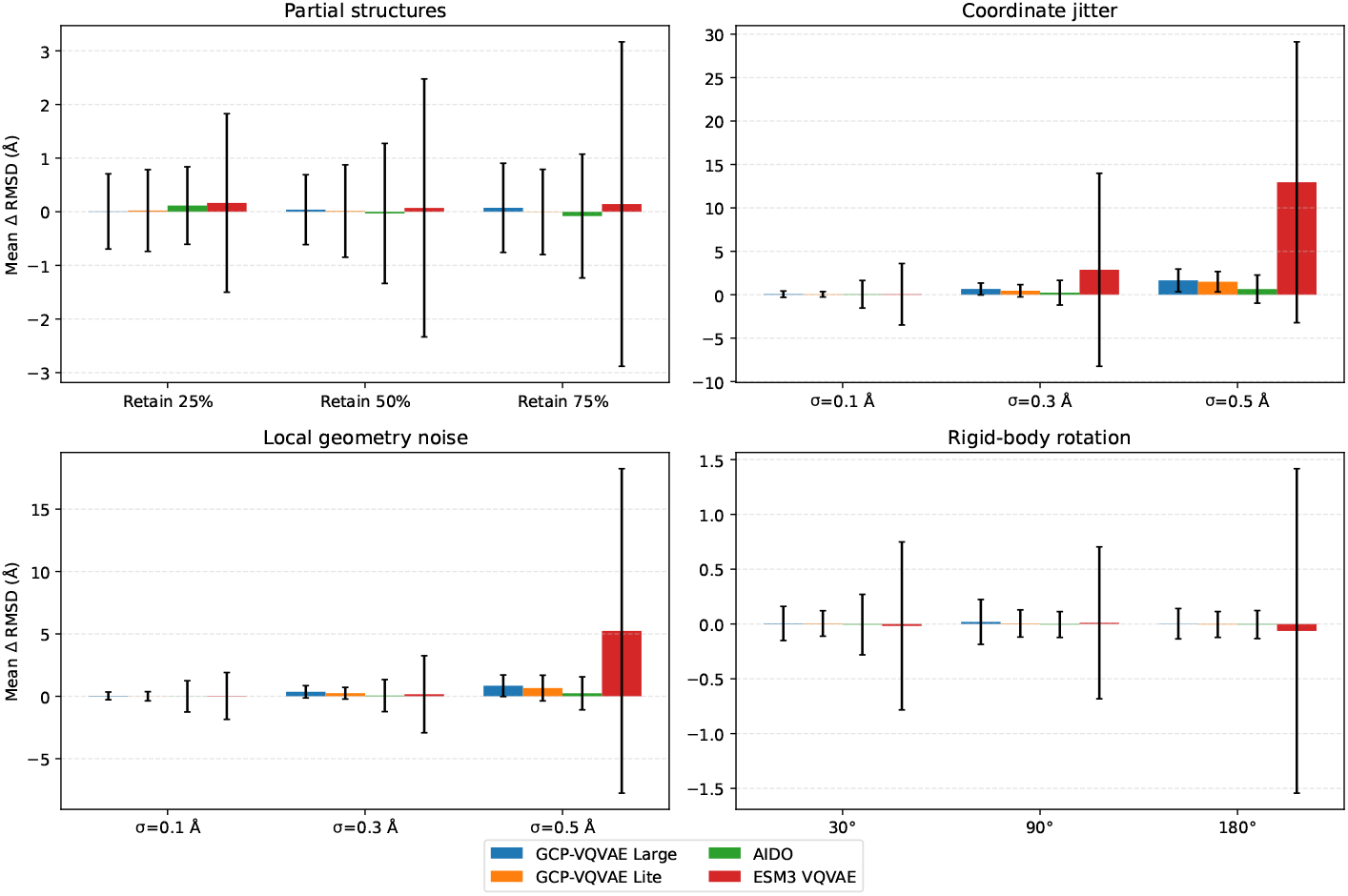
Reconstruction robustness to structural noise and input corruption. We report mean std *±* ΔRMSD (in Å) over 500 zero-shot proteins, where ΔRMSD = RMSD(noisy reconstruction, clean input) – RMSD (clean reconstruction, clean input), computed on C_*α*_ atoms after Kabsch alignment. We evaluate four perturbation families: (top-left) partial structures via contiguous truncation (retain 25/50/75% of residues), (top-right) global coordinate jitter with Gaussian noise 𝒩 (0, *σ*^2^) applied to all backbone atoms, (bottom-left) local geometry noise by perturbing N/C/O around fixed C_*α*_ positions, and (bottom-right) rigid-body rotations of the full backbone. Lower values indicate higher robustness, and rigid-body rotations act as a sanity check where degradation should remain near zero under pose-invariant tokenization.

We set the coefficients *λ* (Equation 10) by monitoring the per-term gradient norms and equalizing their contribution to shared parameters. This gradient-norm balancing prevents any single term (e.g., direction or distance) from dominating updates and yields stable, faster convergence.

Following the structure-tokenizer configuration in ESM-3 (Hayes et al., 2025), we adopt a 4 096-token codebook and select the code embedding dimension to balance reconstruction fidelity and utilization. In particular, the Lite model uses a 128-dim codebook embedding, while the Large model increases this to 256 to better utilize the larger decoder capacity and improve representational bandwidth through the discrete bottleneck. In preliminary sweeps, increasing the code embedding dimension beyond 256 tended to reduce effective codebook utilization, so we use 256 as a practical upper bound in our setting.

Throughout all experiments, sequences longer than the model’s input window are truncated to the first *L* residues (Stage 1: *L* = 512, Stage 2: *L* = 1,280), and all metrics are computed on the truncated region.

#### C.1 Zero-Shot Performance

#### C.2 Inference Speed Benchmarking

We benchmark inference latency for our GCP-VQVAE variants (Large and Lite) alongside top-performing open baselines, ESM3 VQVAE and AIDO StructureTokenizer, across sequence lengths ranging from 32 to 1024 residues. We report the mean wall-clock runtime per protein (in milliseconds) for (i) the encoding pass (backbone coordinates → discrete tokens), (ii) the decoding pass (tokens → reconstructed coordinates), and (iii) end-to-end inference. Benchmarks are conducted on a single Nvidia A6000 Ada 48GB GPU via bf16, using maximum feasible batch sizes (128 for GCP and ESM3; 4 for AIDO, reduced to 1 at *L* = 1024 due to VRAM constraints). Table 11 summarizes the resulting compute profiles, revealing distinct scaling behaviors: While ESM3 is competitive for short sequences, GCP-VQVAE Lite is the fastest method across *all* evaluated sequence lengths, achieving up to ∼ 6 × lower end-to-end latency than ESM3 and ∼ 500 × lower latency than AIDO at 1024 residues.

**Table 11:** Inference latency (ms) for structure tokenization (encoder) and reconstruction (decoder) across different sequence lengths. Total denotes end-to-end time (encode + decode) per protein.

| Seq. Len. | GCP-VQVAE Large |  |  | GCP-VQVAE Lite |  |  | ESM3 VQVAE |  |  | AIDO |  |  |
| --- | --- | --- | --- | --- | --- | --- | --- | --- | --- | --- | --- | --- |
|  | Enc ↓ | Dec ↓ | Total ↓ | Enc ↓ | Dec ↓ | Total ↓ | Enc ↓ | Dec ↓ | Total ↓ | Enc ↓ | Dec ↓ | Total ↓ |
| 32 | 15.53 | 0.17 | <u>15.70</u> | 15.21 | 0.10 | **15.31** | 3.98 | 21.70 | 25.68 | 7.35 | 39.34 | 46.68 |
| 64 | 15.93 | 0.45 | <u>16.38</u> | 15.46 | 0.20 | **15.67** | 4.90 | 21.81 | 26.71 | 6.89 | 38.29 | 45.18 |
| 128 | 16.51 | 1.20 | <u>17.71</u> | 15.79 | 0.69 | **16.48** | 8.60 | 21.87 | 30.46 | 7.37 | 73.22 | 80.59 |
| 256 | 17.18 | 2.63 | <u>19.82</u> | 16.39 | 1.69 | **18.08** | 16.92 | 25.54 | 42.46 | 16.57 | 409.51 | 426.08 |
| 512 | 19.60 | 5.34 | <u>24.93</u> | 17.78 | 3.51 | **21.28** | 34.16 | 52.36 | 86.52 | 35.01 | 2240.84 | 2275.86 |
| 1024 | 25.15 | 12.47 | <u>37.63</u> | 21.46 | 7.52 | **28.99** | 70.96 | 112.40 | 183.36 | 65.52 | 15303.70 | 15369.22 |

**Table 12:**
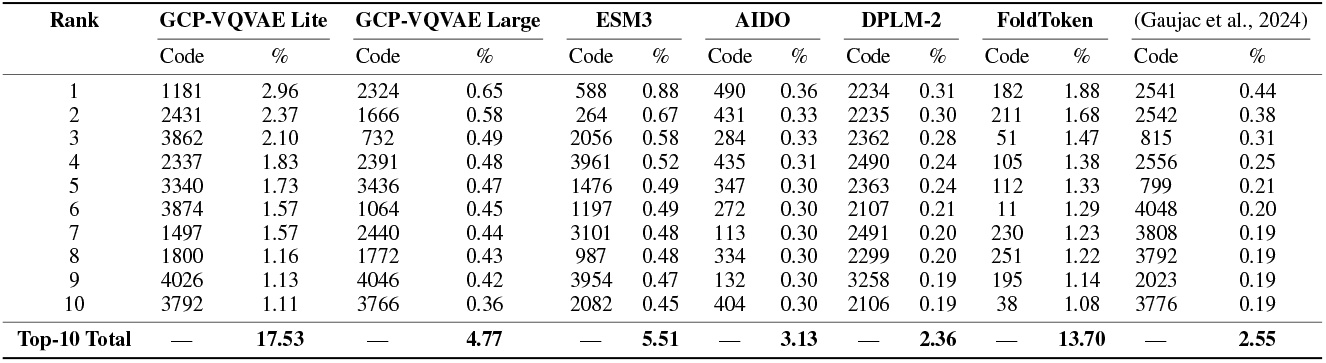
Top-10 most frequent codebook tokens and their usage percentages on the zero-shot test set. We omit the AminoAseed report due to poor reconstruction results.

#### C.3 Reconstruction Robustness to Structural Noise

To assess robustness beyond NaN-masking, we stress-test reconstruction under controlled perturbations of the input backbone coordinates (N, C_*α*_, C, O). For each structure, we first compute a clean baseline reconstruction, then apply a noise transformation to the input coordinates, re-tokenize and reconstruct, and measure the reconstruction degradation as ΔRMSD relative to the clean baseline. Concretely, we report ΔRMSD = RMSD (noisy reconstruction, clean input) – RMSD (clean reconstruction, clean input), where RMSD is computed on C_*α*_ atoms after Kabsch alignment.

We evaluate four perturbation families: (i) partial-structure truncation by keeping only a contiguous segment of the chain, (ii) global coordinate jitter via i.i.d. Gaussian noise applied to all backbone atoms, (iii) local geometry noise by jittering N/C/O around fixed C_*α*_ positions, and (iv) rigid-body rotations. Experiments are run on 500 randomly sampled proteins from our zero-shot PDB 2024– 2025 set, truncated to 1024 residues prior to noise injection, and in figure 16, we report mean std ± ΔRMSD across samples for GCP-VQVAE (Large/Lite) and representative open baselines (ESM3 VQVAE, AIDO).

#### C.4 Representation Evaluation

We note that the remote homology tasks are omitted from the reported results due to data and label missing and inconsistencies. Integrating our custom GCP-VQVAE architecture with the official PST benchmark codebase (Yuan et al., 2025b) was not a straightforward task, as the original framework does not natively support external VQ-VAE models without significant modification. Consequently, we were required to re-implement the evaluation pipeline for each specific task. This extensive reengineering introduced potential uncertainties regarding the alignment between our embeddings and the regression targets.

#### C.5 Codebook Evaluation

To investigate the relationship between learned VQ codebook tokens and protein backbone geometry, we performed a Ramachandran plot analysis on a zero-shot test set comprising 1,938 protein structures not seen during training. For each protein, we passed the original backbone coordinates through the GCPNet encoder and vector quantizer to obtain per-residue VQ code assignments, then decoded these quantized representations to reconstruct the backbone coordinates. We computed backbone dihedral angles *ϕ* and *ψ* for both original and reconstructed structures using standard geometric formulas based on the N, C*α*, and C atomic positions of consecutive residues.

Each residue was plotted in Ramachandran space and colored according to its assigned VQ code index. For visualization clarity, we highlighted the top-30 most frequent codes with distinct colors while displaying remaining codes in gray. We included reference boxes indicating canonical secondary structure regions: the *α*-helix region (*ϕ* ∈ [− 90, − 30], *ψ* ∈ [− 70, − 15]) and the *β*-sheet region (*ϕ* ∈ [− 160, − 80], *ψ* ∈ [− 90, − 180]). These boundaries represent simplified rectangular approximations of sterically allowed regions based on standard Ramachandran conventions.

We analyzed both the Lite and Large model variants across a total of 447,784 residue-level data points. The comparison between original and reconstructed structure distributions allows us to as-sess whether the quantization bottleneck preserves local backbone geometry, while the comparison between model variants reveals differences in codebook utilization and conformational specificity.

To quantitatively assess the information content and utilization of the learned VQ codebook, we computed the following information-theoretic metrics on the discrete token sequences *Z* = (*z*_1_, *z*_2_, …, *z*_*L*_) generated by the encoder:

##### Unigram Statistics

We measure codebook utilization using unigram entropy *H*_1_ = − *p*(*z*) log_2_ *p*(*z*) and perplexity 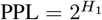. Perplexity represents the effective vocabulary size; an ideal uniform codebook would have PPL = |*K*| = 4096 (*H*_1_ = 12 bits). We report *Effective Usage* as PPL*/*|*K*|.

##### Zipf Distribution Fit

Natural languages typically follow Zipf’s law, where the frequency of the *r*-th most common token scales as *f* (*r*) ∝ *r*^−*b*^. We fit a linear regression to the log-log rank-frequency plot and report the slope *b* and coefficient of determination *R*^2^. A slope near − 1 with high *R*^2^ indicates natural language-like power-law scaling.

##### Conditional Statistics

To evaluate sequential structure, we compute conditional entropies *H*(*Z*_*t*_ | *Z*_*t*−1_) (bigram) and *H*(*Z*_*t*_ | *Z*_*t*−2_, *Z*_*t*−1_) (trigram). Lower conditional entropy indicates higher predictability given context. We report the *Trigram Perplexity* (Tri 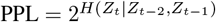), which quantifies the effective branching factor of the sequence given a two-token history.

##### Mutual Information

We measure long-range dependencies using mutual information *I*(*Z*_*t*_; *Z*_*t*+Δ_) = *H*(*Z*_*t*_) − *H*(*Z*_*t*_ | *Z*_*t*+Δ_) at various lags Δ ∈ {1, 4}. Higher mutual information indicates that tokens carry significant predictive information about future states in the sequence.

#### C.6 Major Issues in Performance Analysis

During our evaluation pipeline development, we identified a critical issue affecting TM-score calculations for experimental PDB structures. Unlike computationally predicted structures that typically use sequential residue numbering starting from 1, experimental PDB files from sources such as CASP and CAMEO often contain non-standard residue indices reflecting the original crystallographic or NMR data (e.g., residues numbered 23–97 instead of 1–75). Additionally, experimental structures frequently contain missing residues due to unresolved electron density or flexible loop regions, creating gaps in the residue numbering sequence (e.g., residues 1–45, 52–120) that further complicate alignment. In the first release of the manuscript, when VQ-VAE reconstruction scripts saved decoded structures with default 1-indexed sequential residue numbering while comparing against original PDBs with their native non-sequential numbering, the TM-align algorithm failed to correctly pair corresponding residues, resulting in artificially low TM-scores despite near-identical atomic coordinates. For instance, a structure with actual C*α*-RMSD of 0.30 Å was incorrectly reported as having TM-score of 0.10 due to this mismatch. We corrected this by preserving the original residue indices when writing reconstructed PDB files and applying Kabsch alignment before evaluation, ensuring consistent residue numbering between original and reconstructed structures for accurate metric computation.

### D Compression calculation

Each residue is encoded by one code from a 4 096-entry book, i.e., log_2_(4096) = 12 bits = 1.5 bytes/residue. For *L* = 512 residues, the token stream is 512 × 1.5 = 768 bytes. Raw backbone coordinates (N, C_*α*_, C) stored as 32-bit floats require 3 atoms × 3 coords × 4 bytes = 36 bytes/residue, i.e., 36 × 512 = 18,432 bytes. If we include a tiny global pose header (e.g., rotation+translation) of ≈ 36 bytes, the coded footprint is 768+36 = 804 bytes. Thus the compression ratio is 18 432*/*804 ≈ 22.9 × (approximately 24 × if the pose header is omitted). This estimate concerns backbone only; metadata and format containerization add negligible overhead relative to the raw-float baseline.

### E LLM Usage

In this manuscript, we used large language models only for copy-editing: improving grammar, clarity, and style of author-written text.

## Footnotes

1 March 2024 release.

2 We built the VQVAE components extensively by using the x-transformers and vector-quantize-pytorch libraries.

